# Repertoire analyses reveal TCR sequence features that influence T cell fate

**DOI:** 10.1101/2021.06.23.449653

**Authors:** Kaitlyn A. Lagattuta, Joyce B. Kang, Aparna Nathan, Kristen E. Pauken, Anna Helena Jonsson, Deepak A. Rao, Kazuyoshi Ishigaki, Soumya Raychaudhuri

## Abstract

T cells acquire a regulatory phenotype when their T cell receptors (TCRs) experience an intermediate-high affinity interaction with a self-peptide presented on MHC. Using TCR sequences from FACS-sorted human cells, we identified TCR features that shape affinity to these self-peptide-MHC complexes, finding that 1) CDR3β hydrophobicity and 2) certain TRBV genes promote Treg fate. We developed a scoring system for TCR-intrinsic regulatory potential (TiRP) and found that within the tumor microenvironment clones exhibiting Treg-Tconv plasticity had higher TiRP than expanded clones maintaining the Tconv phenotype. To elucidate drivers of these predictive TCR features, we examined the two elements of the Treg TCR ligand separately: the self-peptide via murine Tregs, and the human MHC II molecule via human memory Tconvs. These analyses revealed that CDR3β hydrophobicity promotes reactivity to self-peptides, while *TRBV* gene usage shapes the TCR’s general propensity for MHC II-restricted activation.

## INTRODUCTION

During T cell development, regulatory T cells (Tregs) acquire their suppressive phenotype when the affinity of their TCR to the peptide-MHC complex (pMHC) is intermediate-high. In most cases, randomly rearranged V, D, and J genes produce a TCR without enough affinity to pMHC, and so most developing T cells do not survive positive selection in the thymus (“death by neglect”). On the other hand, TCRs with too strong affinity to pMHC result in apoptosis and negative selection for the expressing T cell. For the T cells that survive both positive and negative selection, however, a divergence in phenotype emerges: those whose TCRs have lower affinity to pMHC tend to become conventional T cells (Tconvs) and those whose TCRs have higher affinity tend to gain the Treg phenotype^1–8^. Following thymic selection, a crucial prerequisite for the peripheral induction of Tregs is suprathreshold affinity to pMHC, though other factors such as costimulatory signals exert additional influence^7,9^.

The body of evidence that regulatory versus conventional T cell phenotypes are largely driven by TCR signal strength suggests that the developmental fate of CD4^+^ T cells may be influenced by sequence features of the TCR. Indeed, the degree of overlap in TCR sequence between Tregs and Tconvs is minimal compared to T cell samples of the same phenotype^10^. The distinguishing features of Treg and Tconv TCRs could shed light on the determinants of TCR strength, but the majority of extant work has focused on exact sequence matching rather than generalizable TCR sequence features.

To identify all sequence features that influence TCR strength, we examined 5.7×10^7^ TCRβ chain sequences from 65 donors obtained from 6 public data sets (**Table 1**). First, we derived a comprehensive collection of TCR features (**Supplementary Table 1**) and tested them for differential abundance between Tregs and Tconvs in two human cohorts of TCRβ chains from FACS-sorted T cells^11,12^ (**Figure 1a**). From these results, we developed a Treg-propensity scoring system for the TCR (referred to as TCR-intrinsic regulatory potential or TiRP) (**Figure 1b**). Upon confirming its accuracy in two independent cohorts of T cells sampled from the tumor microenvironment, we used TiRP to examine Treg-Tconv plasticity of tumor-infiltrating clones (**Figure 1b-c**). Finally, to shed light on the etiology of the observed TCR sequence biases, we separately examined the two elements of the Treg TCR ligand: 1) the self-peptide and 2) the human MHC II molecule by calculating TiRP in 1) murine Tregs and 2) human memory Tconvs (**Figure 1d**). Our work reveals that CDR3β hydrophobicity promotes reactivity to self-peptides, while *TRBV* gene usage shapes the TCR’s general propensity for activation in the context of human MHC II restriction.

**Table 1.**
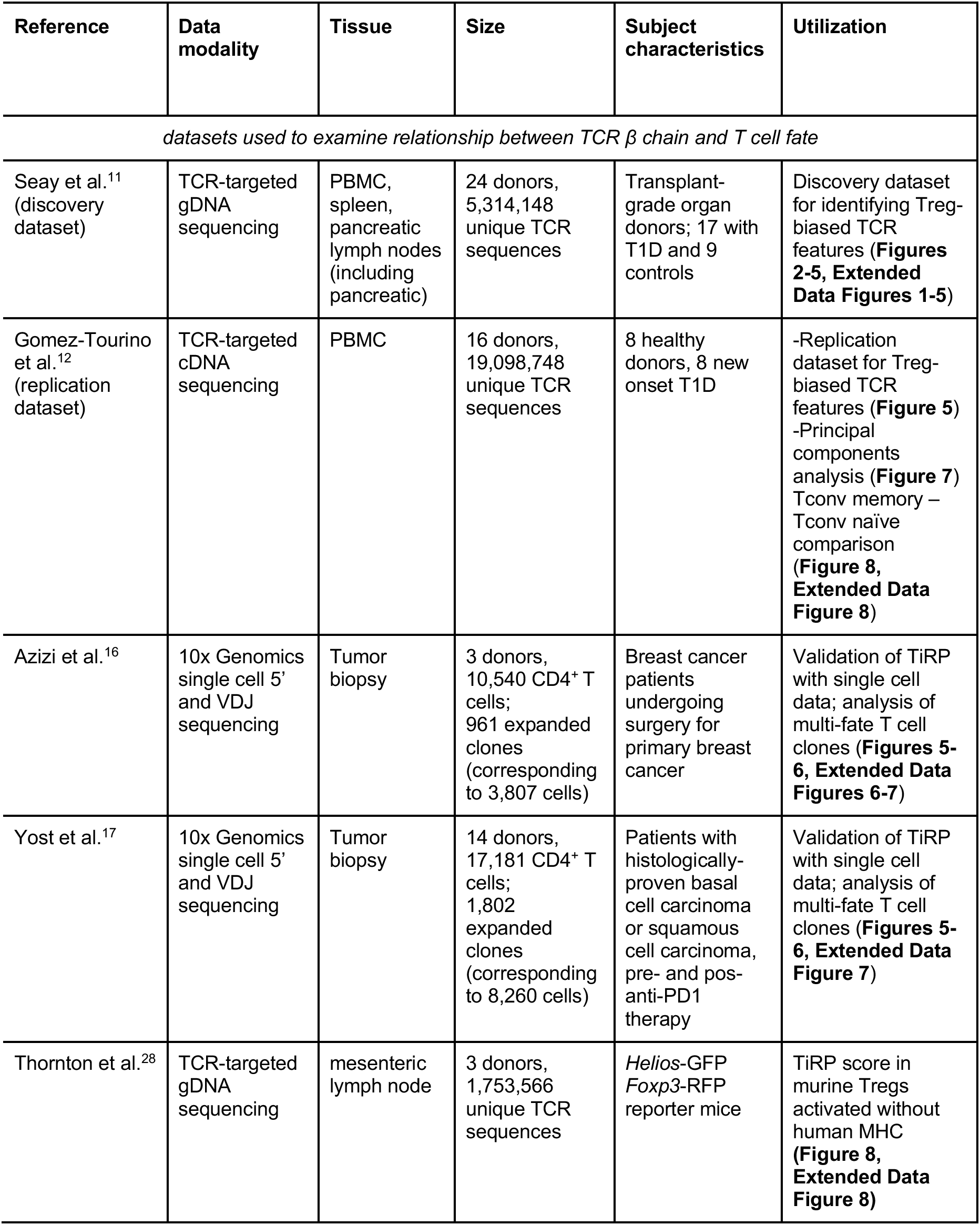
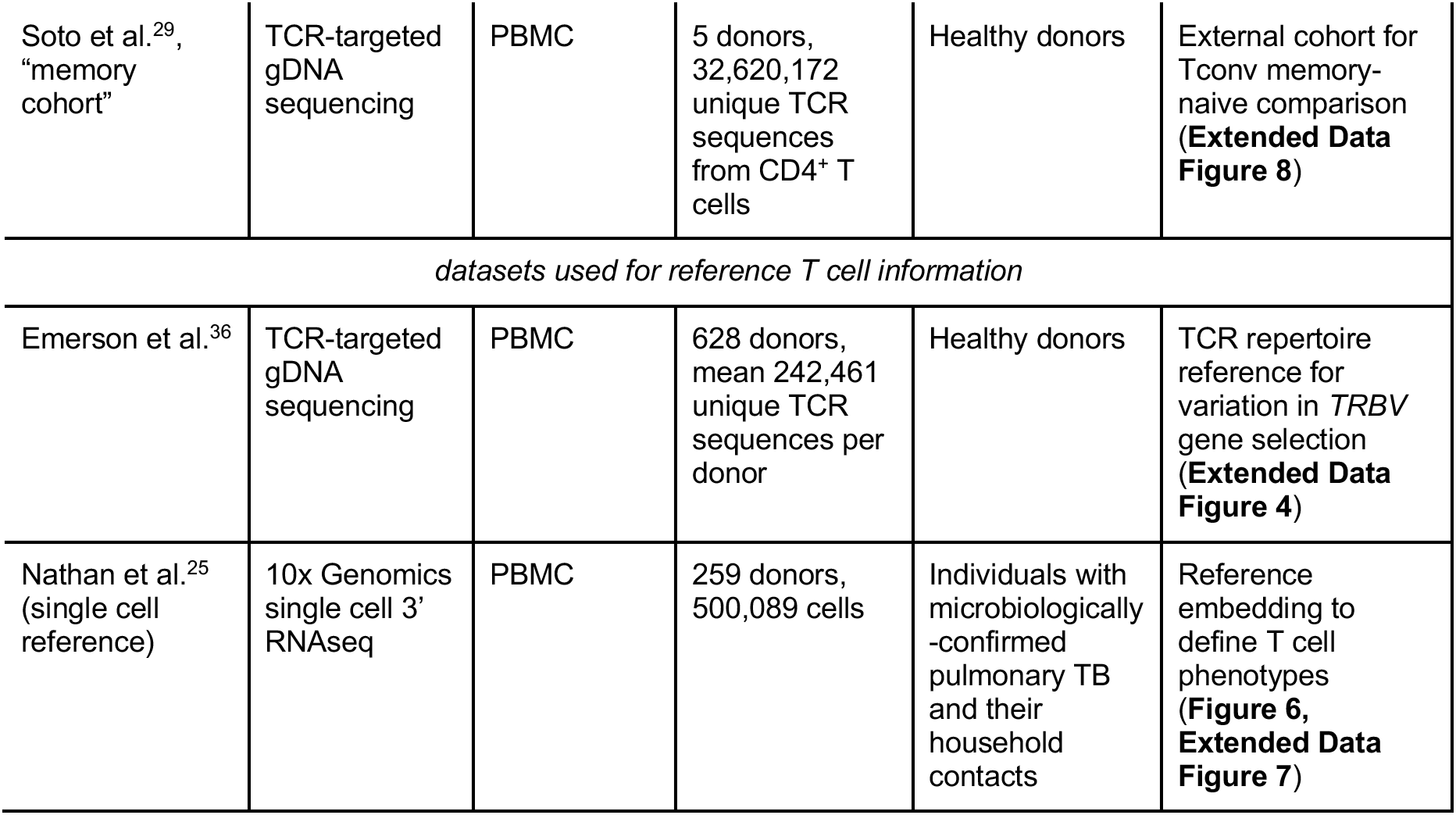
Datasets used in this study.

**Figure 1.**
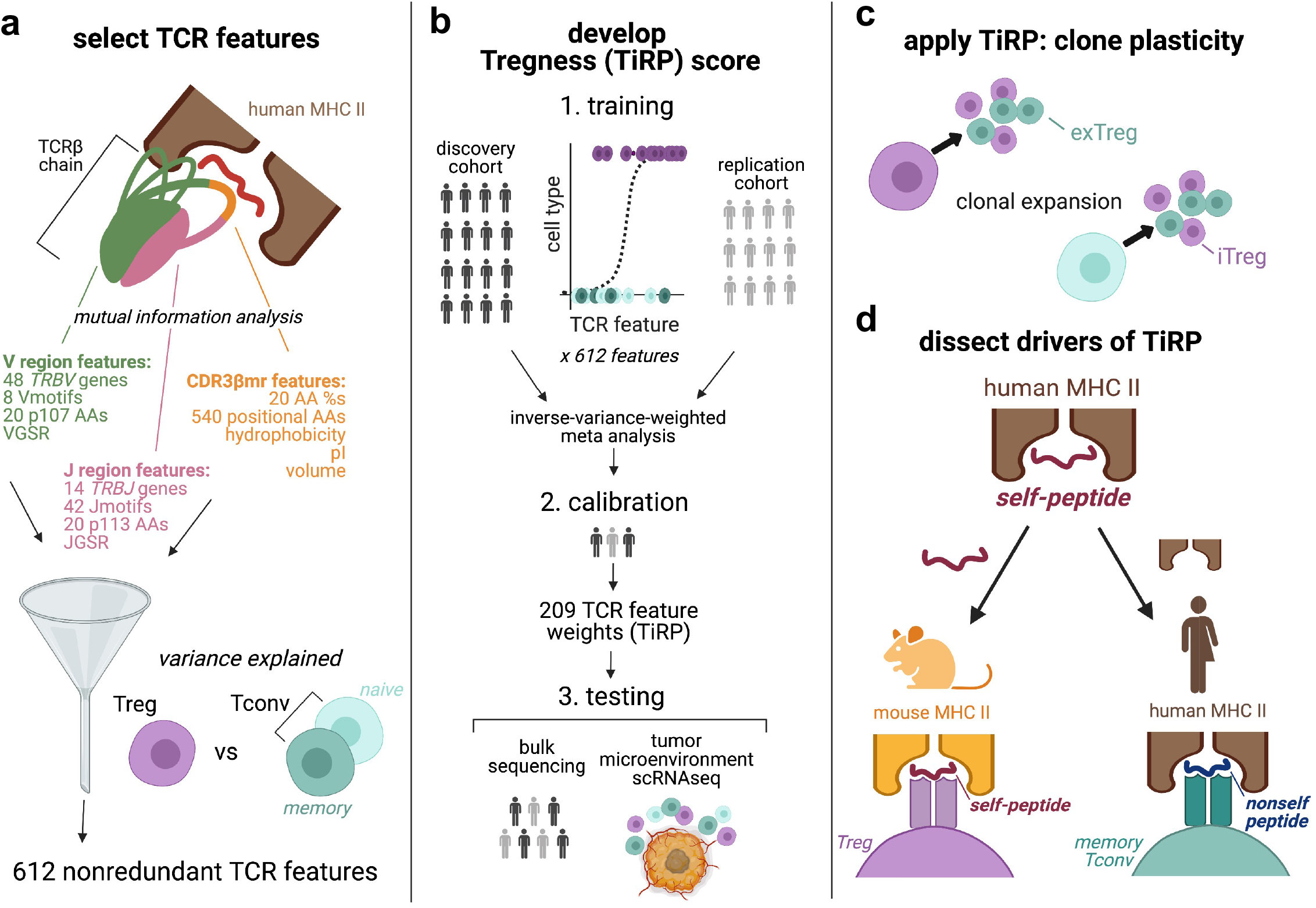
Study design. **(a)** T cell receptor (TCR) β chain in complex with antigenic peptide (red) and human MHC II molecules (brown). The TCR is colored by region: V-region (including CDR1β and CDR2β loops) in green, CDR3β middle region (CDR3βmr) in orange, and J-region in pink. We considered 798 candidate TCR sequence features (Supplementary Table 1) and selected 612 nonredundant TCR features that best explained variance in T cell state (Methods). **(b)** To develop a TCR-intrinsic regulatory potential (TiRP) scoring system, we first split the discovery and replication cohorts into data for training, calibration, and testing. Each human figure represents an individuals’ TCR repertoire sample and is colored according to cohort. We fit logistic regression models for the discovery and replication cohorts separately, and combined the effect sizes for each TCR feature across the two cohorts via inverse-variance-weighted meta-analysis (Methods). We calibrated the *P* value threshold for including a TCR feature in TiRP based on held-out data from both cohorts (Methods). We then tested TiRP in held-out donors from both cohorts, as well as two independent cohorts of T cells sampled from the tumor microenvironment. **(c)** We then examined TiRP in mixed clones: groups of Tregs and Tconvs with the same *TRB* and *TRA* sequences observed in the same individual. These mixed clones likely represent lineages of T cells that have undergone peripheral conversions between the regulatory and conventional phenotypes, which include induced or iTregs (Tconv cells that have acquired a regulatory phenotype) and exTregs (Treg cells that have lost the regulatory phenotype)**. (d)** We then investigated the drivers of TiRP by separately examining the two elements of the human Treg TCR ligand: the self-peptide and the human MHC II molecule. Vmotif: IMGT position 104 – 106 tripeptide, AAs: amino acids, VGSR: V gene selection rate (Supplementary Note), pI: isoelectric point, Jmotif: IMGT position 114-118 sequence, JGSR: J gene selection rate (Supplementary Note). Figure created with Biorender.

## RESULTS

### Defining independent features of the T cell receptor sequence

The TCR is a membrane-anchored heterodimeric protein consisting of an α and a β chain. Each of the two chains includes three highly variable peptide loops that protrude into the TCR-pMHC complex. The most variable of these loops is the CDR3β region in the β chain which mediates recognition of specific antigens. Because *TRBV, TRBD,* and *TRBJ* genes each encode regions of CDR3β, we anticipated that the CDR3β sequence would feature blocks of strongly correlated residues. To determine the boundaries of these correlated regions, we examined the mutual information structure of CDR3β peptides in a previously published cohort of targeted TCR sequencing in multiple tissues and PBMCs^11^ (“discovery cohort”, **Table 1**). To assess generalizability of any findings, we held out data from six randomly selected donors (**Figure 1b, Methods**).

Pairwise mutual information calculations between CDR loop residues revealed three distinct regions of the TCR: the V-region (IMGT position 1–107), CDR3β middle region (CDR3βmr, p108–p112), and J-region (p113–p118) (**Figure 2a-b**, **Extended Data Figure 1**). While random nucleotide insertions in the highly variable CDR3βmr obscured the identity of the *TRBD* gene, the germline-encoded V- and J- regions demonstrated sequence conservation and high inter-residue mutual information (**Figure 2a**). Mutual information was concentrated at the flanking ends of CDR3β such that eight p104-p106 tripeptides (“Vmotifs”) and 42 p113-p118 pentapeptides (“Jmotifs”) accounted for >90% of observations. Upon observing minimal mutual information between the three regions, we elected to undertake a three-pronged modeling approach, in which we would examine the V-, middle, and J- regions independently.

**Figure 2.**
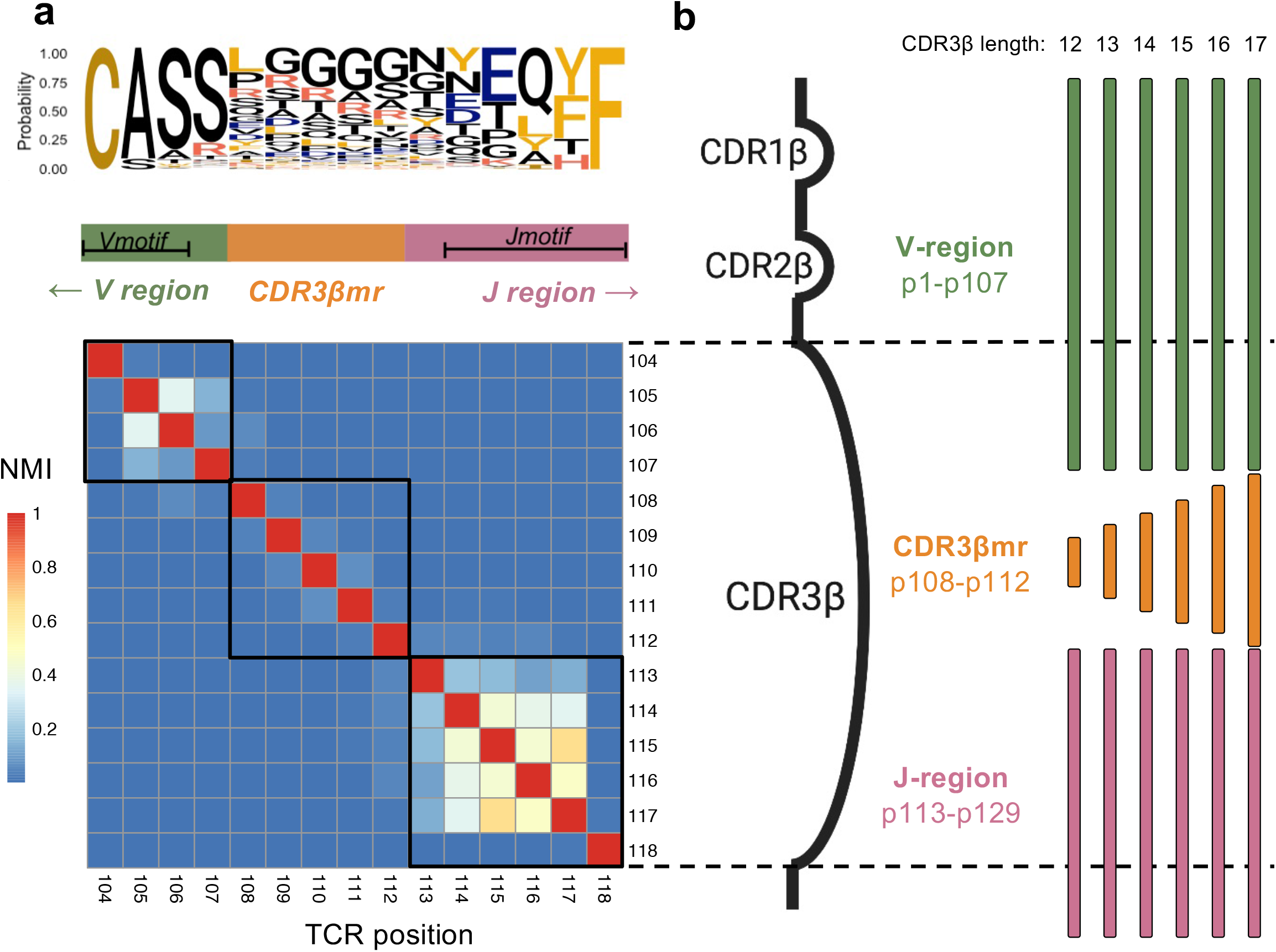
TCR sequence structure. **(a)** Probability of each amino acid in each CDR3β position depicted by a sequence logo, with a heatmap of normalized mutual information (NMI) between each pair of CDR3β residues for the most frequent CDR3β length, 15 amino acids. Based on this mutual information structure, we partitioned the CDR3β sequence into a Vmotif within a V-region, a CDR3β middle region (CDR3βmr), and a Jmotif within a J-region. **(b)** Schematic showing TCRs of multiple lengths aligned to the TCR β chain structure. Three complementary-determining regions within the TCR β chain protrude as loops into the pMHC-TCR complex: CDR1β, CDR2β, and CDR3β. CDR1β and CDR2β are encoded by the *TRBV* gene, while CDR3β spans *TRBV*-encoded residues, random nucleotide insertions (CDR3βmr) and *TRBJ*-encoded residues. Random nucleotide insertions from VDJ recombination occur at the V/D and D/J junctions, creating variation in CDR3βmr length. Regions suggested by mutual information structure are not drawn to scale. NMI: Normalized mutual information

### Regulatory T cells use specific amino acids within the CDR3β middle region

We first examined the middle region of CDR3β (“CDR3βmr”) of Tregs (CD4^+^CD127^−^ CD25^+^) and Tconvs (CD4^+^CD127^+^) in the discovery cohort. Calculating the mean percentage of CDR3βmr residues occupied by each amino acid yielded strikingly consistent Treg-Tconv differences across donors: Phenylalanine (F), Leucine (L), Tryptophan (W), and Tyrosine (Y) were consistently enriched in Tregs, while Aspartic acid (D) and Glutamic acid (E) were consistently enriched in Tconvs (**Figure 3a**, **Extended Data Figure 2a**). Categorization of amino acids by physicochemical features showed that hydrophobic amino acids were enriched in Tregs, while negatively charged amino acids were enriched in Tconvs (**Extended Data Figure 2b**).

**Figure 3.**
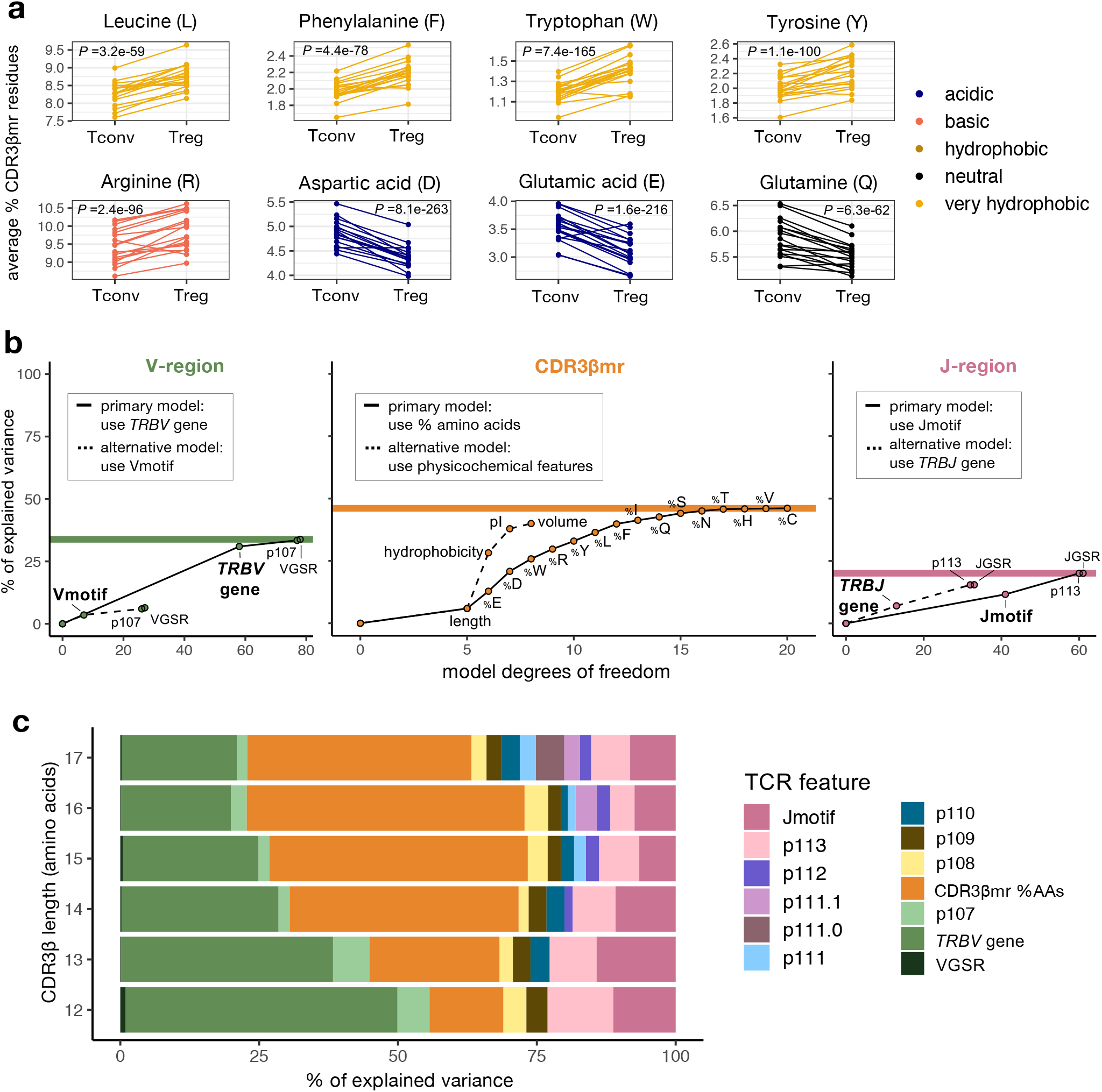
Broad differences exist between the TCRs of Tregs and Tconvs. **(a)** Percentage of select amino acids in the CDR3βmr, plotted as the mean for each donor sample in the discovery cohort, separated by cell type and colored by amino acid groups. *P* values are computed by a Wald test on the coefficient for each amino acid term in a mixed effect logistic regression model (Methods). **(b)** Incremental variance explained by the addition of labeled TCR features to the V-region (left), CDR3βmr (middle), and J-region (right) mixed effect logistic regression models. For each region, the primary modeling approach was compared to the alternative modeling approach, and the modeling approach that explained greater variance was selected. Colored horizontal lines depict the total percent of explained variance attributable to each TCR region, summing to 100%. **(c)** Percent of explained variance by each TCR feature type, summing to 100% for each length of CDR3β. VGSR = V gene selection rate (Supplementary Note). CDR3βmr %AAs = percent composition of amino acids in the CDR3βmr.

To statistically assess these differences, we used nested conditional mixed effect logistic regression models which account for inter-individual differences such as those driven by HLA genotype and tissue source (**Methods**). We observed that 15 amino acids had an independent effect on Treg fate (**Supplementary Table 2, Methods**). To confirm that these effects were consistent across donors and clinical phenotypes, we estimated them in each of the 18 individuals and in T1D cases and controls separately. We found consistent effect sizes in all contexts (**Extended Data Figure 3a-b, Supplementary Table 2, Methods**). We compared this model to an alternative approach in which CDR3βmr was scored by physicochemical features (hydrophobicity, isoelectric point (pI), and volume) rather than percentages of individual amino acid residues (**Supplementary Table 3**, **Methods**). Physicochemical features did not capture as much information as amino acid percentages (**Figure 3b**, middle); hence, we proceeded with an amino acid-based model of the CDR3βmr.

We then ran a separate mixed effects model for each CDR3βmr position (IMGT p108 - 112), testing whether the amino acid at the given position explained variance in T cell fate beyond that accounted for by the CDR3βmr amino acid percentages (**Methods**). We found that each position indeed conveyed additional information regarding the likelihood of Treg fate, but these position-specific effects all together did not explain as much variance as the general amino acid composition of the CDR3βmr (**Figure 3c**, **Supplementary Table 4**).

### CDR3β V and J regions explain variance in T cell state

We then examined the V-region of the TCR. Previous studies have established that genetic variation in the MHC locus shapes the frequency with which *TR(A/B)V* genes are used in the repertoire^13^. Interestingly, MHC polymorphisms explained far more variance in *TRAV* gene usage compared to *TRBV^13^*, consistent with protein structure data demonstrating that *TRAV* contacts MHC at polymorphic sites while *TRBV* contacts MHC at conserved sites^14^. We hypothesized that variation in *TRBV*-encoded residues may alter TCR affinity to these conserved MHC sites, and thereby influence T cell fate.

To test this hypothesis, we extracted sequence features from the V-region and tested their association with Treg fate using mixed effects logistic regression (**Methods**). Through model comparisons, we found that a joint model including *TRBV* gene identity and p107 best represented the region, since the 58 *TRBV* genes explained far more variance than the eight Vmotifs (**Figure 3b** left**, Methods**). To account for inter-individual variation in *TRBV* gene selection, we derived a thymic selection parameter (VGSR) for each *TRBV* gene as a covariate (**Supplementary Note**, **Extended Data Figure 4**). Despite controlling VGSR, *TRBV* gene identity continued to explain a significant amount of variance in T cell fate, with three *TRBV* genes reducing the odds of Treg fate by more than 30% compared to the reference (most common) gene, *TRBV05-01* (*P =* 1.3 × 10^−804^, LRT, **Supplementary Table 5**). As in the CDR3βmr analysis, we confirmed that these associations replicated in models isolated to each individual and to both case and control cohort subsets (**Extended Data Figure 3c-d, Supplementary Table 5**). The consistency in *TRBV* gene effects across individuals suggests that their influence on Treg fate indeed occurs through interactions with conserved MHC residues, and is largely independent of MHC variability between individuals.

We then examined the J-region. In contrast to the V-region, wherein strong p104-p106 sequence conservation constrained multiple *TRBV* genes to the same Vmotif, variable nucleotide editing at the *D/J* junction resulted in multiple Jmotifs associated with each *TRBJ* gene. The 42 Jmotifs explained slightly more variance than the 13 *TRBJ* genes (**Figure 3b,** right), and so we proceeded with a joint model containing the Jmotif and p113 residue. Across all three regions, the most important TCR features for T cell fate determination were the *TRBV* gene identity and the percent composition of amino acids in the CDR3βmr (**Figure 3c**).

### Treg TCRs are enriched for CDR1β apex positive charge and CDR3β middle region hydrophobicity

We wanted to localize physicochemical effects underlying CDR3βmr residue enrichments to specific sequence positions. At each CDR(1-3)β loop amino acid position, we estimated the importance of hydrophobicity, isoelectric point (pI), and volume in influencing Treg fate using a ridge regression model (**Methods, Supplementary Table 6**). Intriguingly, these results provided a physicochemical basis for some of the *TRBV* gene differences observed. Tregs were enriched for positively charged amino acids at p37 of CDR1β (**Figure 4a**). Seven *TRBV* genes assessed in our models harbor a negatively charged residue at p37; all seven of these were significantly depleted for Tregs compared to the reference gene *TRBV05-01*, which has a positively charged Arginine (R) at p37 (**Figure 4b**). As expected from our earlier findings, CDR3βmr featured positive coefficients for hydrophobicity in every position (**Figure 4a**). At each of these positions, every standard deviation increase in hydrophobicity led to a 2.5% (L17, p113) – 6.3% (L12, p113) increase in odds of Treg fate (OR = 1.025, 95% CI = 1.011–1.039, Wald test *P =* 2.7 × 10^−4^ for L17-p113; OR = 1.063, 95% CI = 1.051–1.074; Wald test *P =* 5.2 × 10^−28^ for L12-p113, **Extended Data Figure 5, Supplementary Table 6**).

**Figure 4.**
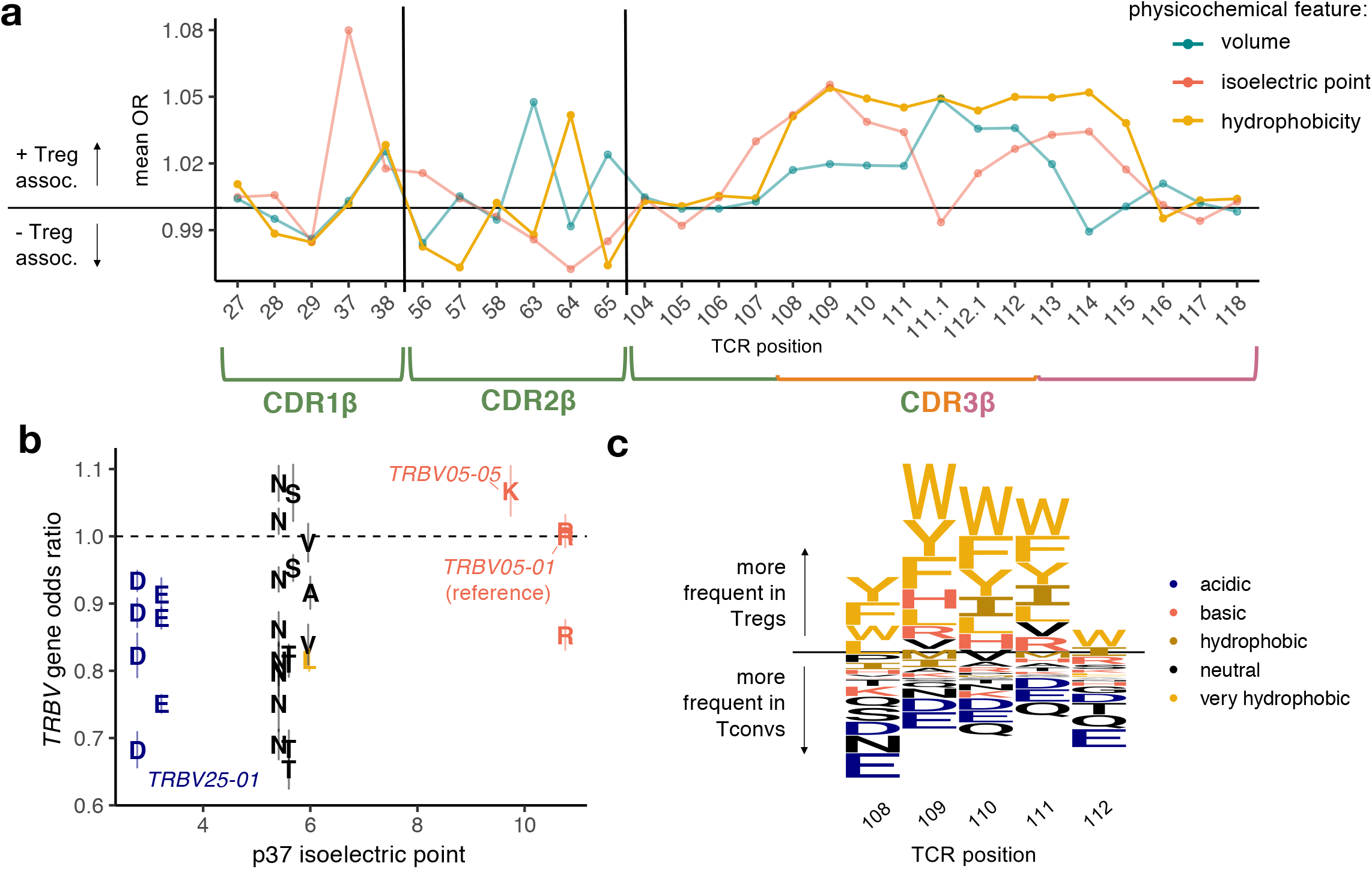
Tregs exhibit position-specific TCR sequence features. **(a)** Estimated odds ratio (per standard deviation) for each physicochemical feature at each CDRβ(1-3) loop position; features with an estimate > 1 are positively associated with Treg fate while features with an estimate < 1 are negatively associated. Odds ratios denote the change in Treg odds per standard deviation increase in the given physicochemical feature at the given TCR position. For each CDR3β length, all effects were estimated jointly in an L2-regularized logistic regression with 10-fold cross-validation (Methods). Shown are the odds ratio estimates for each position-feature averaged across the six CDR3β lengths. Vertical lines denote the boundaries of each CDRβ loop. **(b)** Correspondence between *TRBV* gene isoelectric point at p37 (apex of CDR1β) and *TRBV* gene odds ratio for Treg fate compared to the reference gene, *TRBV05-01*. Each *TRBV* gene is labeled with its amino acid residue at p37 and the 95% confidence interval for its odds ratio. **(c)** Sequence logo depicting the effects of amino acids in the highly entropic CDR3βmr residues, sized proportionally to the associated change in Treg odds, with amino acids more frequent in Tregs above the horizontal line and amino acids more frequent in Tconvs below.

To directly visualize the amino acids associated with Treg fate, we generated a sequence logo representation of the CDR3βmr based on differential amino acid usage at each position (**Figure 4c**, **Methods**). Our results are consistent with the recent finding by Stadinksi *et al.* that hydrophobicity at p109 and p110 promotes the development of T cells that recognize self-antigens^15^. Importantly, we show that this principle extends beyond p109-110 throughout the stretch of entropic CDR3β residues.

### Reproducing TCR associations in an independent data set

Having identified TCR features associated with Treg identity, we next sought to validate them in a public dataset of TCRβ sequences from sorted Treg (CD4^+^CD25^high^CD127^low^) and Tconv (CD4^+^CD25^low^CD27^+^) cells sampled from the peripheral blood of 16 donors^12^ (“replication cohort”, **Table 1**). Despite a different distribution of tissue sources in this data set, the CDR3βmr amino acid percentage effects were nearly identical (Pearson *R =* 0.95, *P =* 4.6 × 10^−8^, **Figure 5a**, **Supplementary Table 2**). Effects for individual *TRBV* genes, Jmotifs, and position-specific amino acid effects were also consistent with discovery (Pearson *R =* 0.56, *P =* 7.5 × 10^−57^, **Figure 5b**, **Supplementary Tables 4-5, Methods**). In the replication cohort, *TRB* sequences were collected by reverse transcription and amplification of RNA rather than direct DNA sequencing. Thus, relative changes in Treg likelihood induced by these TCR sequence features are not only robust to different tissue sources, but also to technical differences in sorting and sequencing protocols.

**Figure 5.**
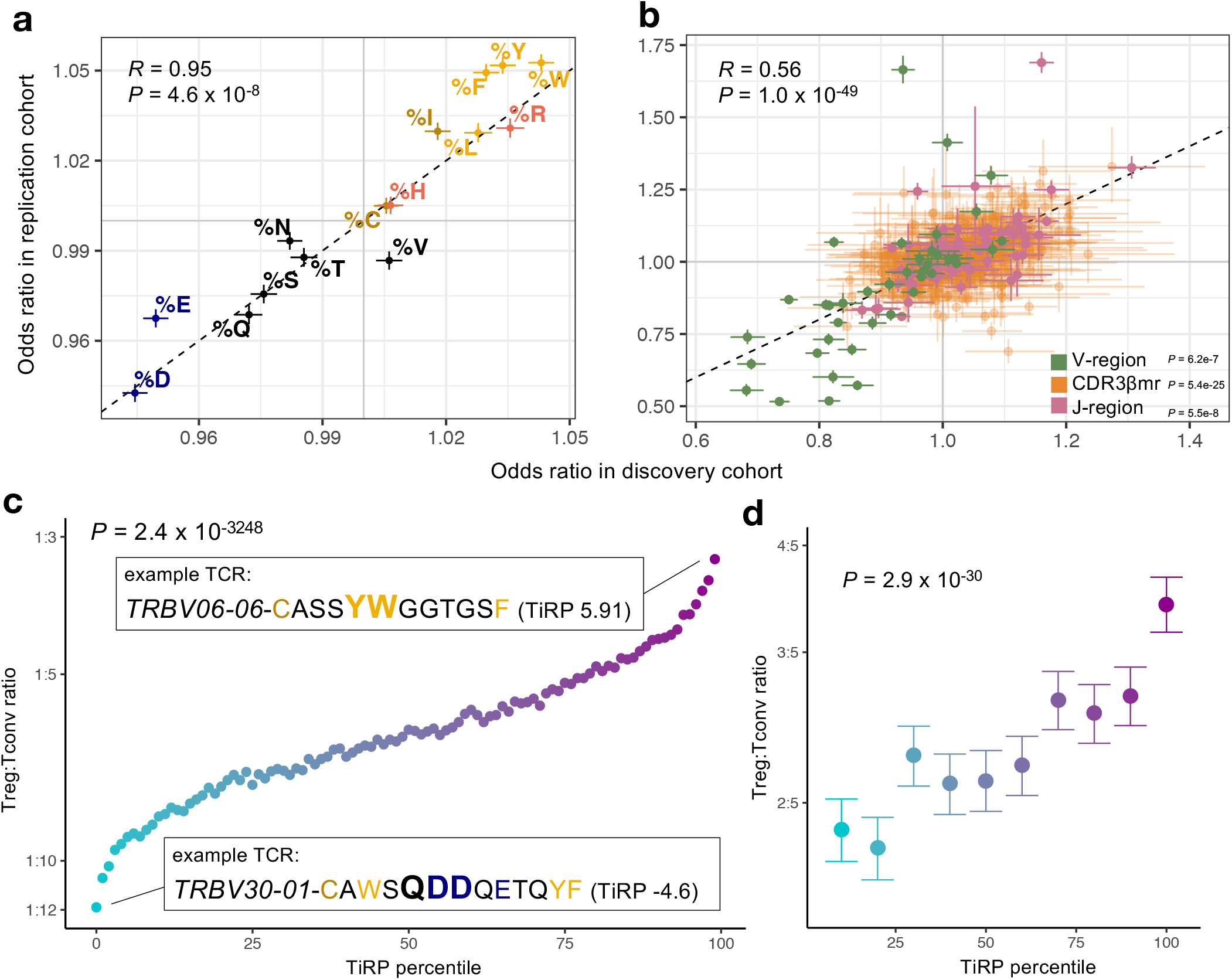
Treg TCR sequence biases replicate in an independent cohort. **(a)-(b)** Correspondence between the discovery and replication cohort odds ratios for all 612 nonredundant TCR features. Odds ratios for CDR3βmr percent composition amino acids are shown on the left; OR corresponds to the change in Treg odds associated with one standard deviation increase in the given CDR3βmr amino acid percentage. Colors for amino acids correspond to the same categorization scheme presented in Extended Data Figure 2b. All other TCR features are shown on the right; OR corresponds to the change in Treg odds associated with the presence of the given TCR sequence feature compared to the reference feature (Supplementary Table 1). **(c)** Validation of TCR-intrinsic regulatory potential (TiRP) score in held-out donors of the discovery and replication dataset (n = 3,277,036 TCRs). Each standard deviation increase in TiRP was associated with a 23% increase in the odds of Treg status (OR: 1.231, 95% CI: 1.227 – 1.235, *P =* 2.4 × 10^−3248^, LRT). Percentile points are colored by Treg:Tconv ratio ranging from blue (lowest) to purple (highest). An example TCR from the lowest and highest percentile are given, with the CDR3βmr highlighted in bold and amino acids colored by the categorizations depicted in Extended Data Figure 2b. **(d)** Validation of TiRP in scRNAseq of CD4+ tumor microenvironment T cells^16,17^ (n = 27,721 cells). Each standard deviation increase in TiRP was associated with an 18% increase in the odds of Treg status (OR: 1.18, 95% CI: 1.15–1.21, LRT *P =* 2.9 × 10^−30^). Error bars outline 95% confidence intervals for Treg/Tconv odds in each TiRP score decile, computed by bootstrap resampling (Methods). *P* value is computed by a LRT between mixed effects logistic regression models differing only in the inclusion of TiRP score as a fixed covariate, one observation per cell (Methods).

### Developing TiRP: a Treg-propensity scoring scheme for the TCR

Having replicated the effect of a comprehensive set of TCR features in two independent cohorts, we next developed a method to quantify the TCR-intrinsic regulatory potential (“TiRP”) of a T cell. Briefly, for a given TCR, TiRP is the sum of Treg association effect sizes of independent sequence features in all three TCR regions (**Methods**). We used meta-analytic effect size estimates across the two cohorts and included only features with a significant effect on T cell fate based on a Bonferroni *P* value threshold (**Methods**). As a result, TiRP is the weighted sum of 25 *TRBV* genes, 23 Jmotifs, 4 CDR3β lengths, 14 CDR3βmr amino acid percentages, and 143 position-specific features (**Supplementary Table 7**).

We then tested our TiRP score on the four discovery cohort donors and two replication cohort donors whose repertoire data had been withheld from all former analyses. We observed that a one standard deviation increase in TiRP in these held-out data resulted in a 23% increase in the odds of Treg status (OR: 1.231, 95% CI: 1.227 – 1.235, LRT *P =* 2.4 × 10^−3248^, **Figure 5c**, **Supplementary Table 8**, **Methods**). TCRs in the highest-scoring decile were more than twice as likely as TCRs in the lowest-scoring decile to belong to a Treg: 1 in every 3.9 compared to 1 in every 9.1. To ensure that this TCR-T cell state covariation was contingent on the biology of surface-expressed TCRs, we repeated this analysis on the nonproductive TCRs in the four held-out donors for which out-of-frame reads were available (**Methods**). This indeed abrogated the association between Tregness score and Treg fate (OR: 1.00, 95% CI: 0.97 – 1.04, LRT *P* =0.96).

### TiRP helps to explain Treg-Tconv plasticity in the tumor microenvironment

To externally validate our scoring system, we aggregated scRNAseq data from 27,721 tumor-infiltrating CD4^+^ T cells with paired TCR reads from two publicly available 5’ scRNAseq datasets^16,17^(**Table 1**). We scored the *TRB* chain of each cell and assessed whether the TiRP score explained variance in T cell phenotype, as defined by the original authors for the Yost et al. cohort and by a standard clustering pipeline for the Azizi et al. cohort (**Methods, Extended Data Figure 6a-b**). Consistent with our previous observations, there was a nearly two-fold increase in Treg likelihood in the top TiRP decile compared to the bottom TiRP decile in these data (**Figure 5d**, OR for top-bottom decile comparison: 1.68, 95% CI: 1.49-1.90, *P =* 2.3 × 10^−17^; OR for all cells: 1.18 per unit increase in TiRP, 95% CI: 1.15–1.21, LRT *P =* 2.9 × 10^−30^, **Supplementary Table 8**).

We next asked whether TiRP could help to explain regulatory T cell plasticity. It is well-recognized that naïve Tconv thymic emigrants can be peripherally induced to adopt a regulatory phenotype^18,19^. Conversely, some Tregs have been observed to lose *FOXP3* expression and adopt a pro-inflammatory phenotype^20–23^ (“exTregs”, **Figure 1c**). Expanded T cell clones (possessing the same TCR) observed as both Tregs and Tconvs within the same donor (hereafter referred to as “mixed clones”) may represent lineages of T cells that have undergone such peripheral conversions. We hypothesized that the TiRP of these T cells may be intermediate, rendering them most susceptible to peripheral conversion.

Before testing our hypothesis, we used Symphony^24^ to standardize cell type definitions across the two cohorts by mapping cells of expanded clones from both datasets (12,067 cells) into a common reference atlas^25^ of T cell states based on joint transcriptional and proteomic profiling (**Figure 6a-c**, **Table 1**, **Extended Data Figure 7a-d**, **Methods**). On average, 19.2% of expanded clones from the same donor were observed in both the Treg and Tconv state, including a few large clones with a relatively even balance (**Figure 6d-e**, **Supplementary Table 9**).

**Figure 6.**
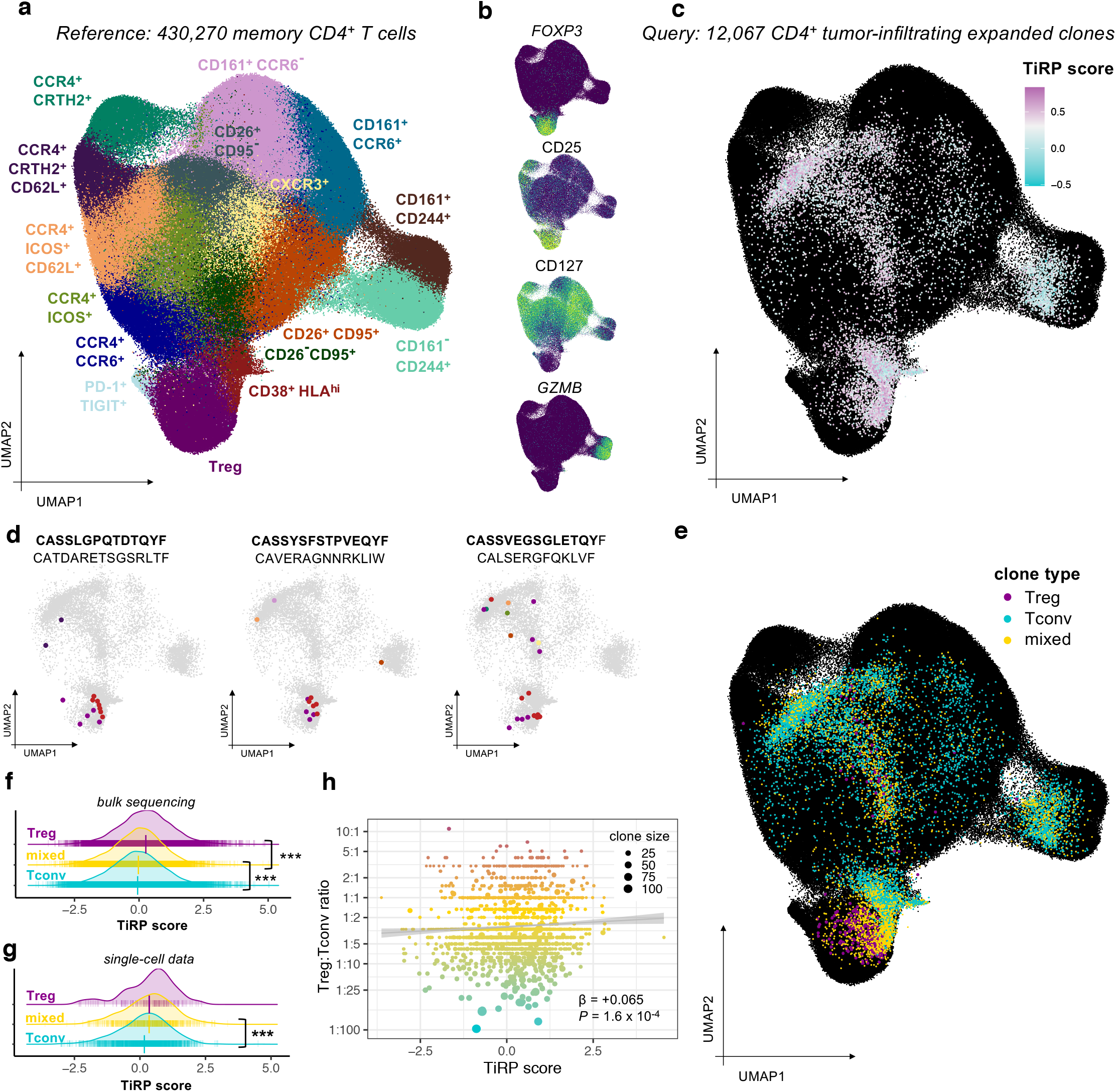
TiRP helps to explain clonal plasticity in the tumor microenvironment. **(a)** Reference T cell dataset, colored by cell type clusters according to transcriptional and surface marker variation depicted in Extended Data Figure 6c-d. **(b)** Select gene expression (*FOXP3, GZMB*) and surface marker abundance (CD25, CD127) for cells in the reference T cell dataset (low = purple, high = light green). **(c)** Tumor microenvironment T cells of expanded clones mapped into the reference embedding by Symphony. Each cell is colored by the TiRP score of its paired *TRB* chain, with KNN smoothing for visualization (Methods). TiRP is scaled such that 0 corresponds to the mean score and one unit corresponds to one standard deviation of held-out bulk sequencing TCRs (Figure 5c). **(d)** Cell members of three example mixed clones are highlighted in color according to their cell type classification by Symphony (colors as in (a)). Within a given plot, each cell expresses the same *CDR3β* DNA sequence, the same CDR3α amino acid sequence, and was found within the same donor (CDR3β amino acid sequence listed above CDR3α amino acid sequence for each plot). **(e)** Same as (b), with each cell colored according to the type of its parent clone: Treg for clones containing only Treg cells, Tconv for clones containing only Tconv cells, and “mixed” for clones containing both Treg and Tconv cells. **(f)** TiRP score distribution of purely Tconv, purely Treg, and “mixed” expanded clones from held-out bulk sequencing data. *P =* 2.0 × 10^−40^ for mixed-Tconv difference, *P =* 9.1 × 10^−16^ for mixed-Treg difference. **(g)** TiRP score distribution of purely Tconv, purely Treg, and mixed expanded clones in tumor-infiltrating scRNAseq data. *P =* 3.0 × 10^−4^ for mixed-Tconv difference, *P =* 0.55 for mixed-Treg difference. For (f) and (g), vertical bars denote mean and standard error of the mean per clone type. **(h)** Correspondence between TiRP score and the composition of T cell states within each clone, quantified by the Treg:Tconv ratio. Best fit line is shown in gray; clones are colored by Treg:Tconv ratio. β corresponds to the slope of the regression line between the log-transform of the Treg:Tconv ratio and TiRP score. *P* value is computed by the LRT between mixed effect logistic regression models (Methods).

We next tested whether the TiRP score of mixed clones was in between that of purely Tconv and Treg clones (**Methods**). In the previously held-out bulk sequencing data, the TiRP scores of mixed clones were significantly greater than those of expanded Tconv clones and less than those of expanded Treg clones (**Figure 6f,** mixed-Tconv difference = 0.03, *P =* 2.0 × 10^−40^; mixed-Treg difference = −0.29, *P =* 9.1 × 10^−16^, LRT, **Methods**). These single cell data confirmed that Tregs of mixed clones indeed exhibited greater *FOXP3* expression than Tconvs within the same clonal expansion (**Extended Data Figure 7e**, **Methods**). As in the previously held-out bulk sequencing data, mixed clones in single cell data had intermediate TiRP scores which were significantly greater than the scores of expanded, pure Tconv clones (**Figure 6g**, mixed-Tconv mean TiRP difference = 0.182, *P =* 3.0 × 10^−4^, LRT, Methods). With the limited extent of Treg expansion, we were underpowered to detect significant differences between mixed and Treg clones in these data (mixed-Treg mean TiRP difference = −0.005, *P =* 0.57, LRT). When we quantified clone phenotypes by the proportion of Tregs and Tconvs within each clone, increasing TiRP corresponded to more Treg-skewed clonal expansions (LRT *P =* 0.003, **Figure 6h**, **Methods**). To our knowledge, TiRP is the first metric to identify TCR-intrinsic, rather than TCR-extrinsic factors relevant to peripheral phenotypic conversion.

### Separable components of TiRP: affinity to self-peptides and affinity to human MHC

We next asked whether TiRP captured the major sources of TCR sequence variation between sorted T cell samples from diverse individuals. For this, we conducted a principal components analysis (PCA) of TCR feature frequencies in the sorted samples of the replication dataset, in which all T cell states of interest were available (**Methods**). We observed that the major axes of TCR sequence variation corresponded to T cell state, rather than donor HLA genotype or clinical phenotype (**Figure 7a**, **Extended Data Figure 8a-b**). While our previous supervised modeling was designed to focus on Treg-Tconv differences, this approach recovered the importance of T cell state in an unsupervised manner.

**Figure 7.**
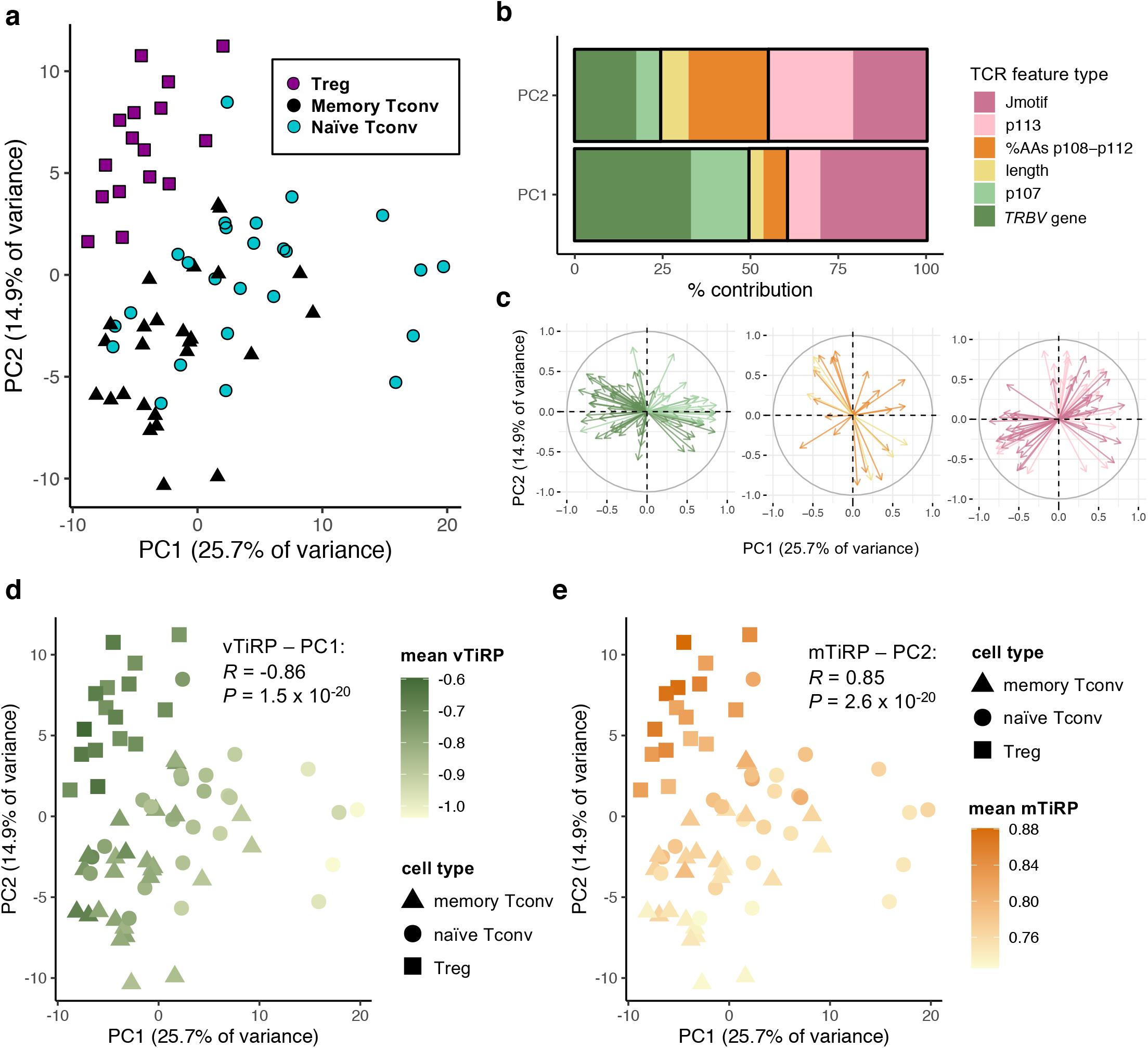
Two axes of TCR-driven cell states. **(a)** 67 samples from the replication cohort colored by cell type and arranged by principal component space according to variation in TCR sequence feature frequencies (Methods). **(b)** Percent contribution of each type of TCR sequence feature to the first two principal components. **(c)** Loadings of each of the TCR sequence features on PC1 and PC2, depicted by arrows, separated by TCR region and colored by the same scheme as in (b). **(d)** Samples arranged in PC space as in (a), colored by mean TiRP in the V-region of the TCR (vTiRP). **(e)** Same as in (d), colored by mean TiRP in the CDR3βmr (mTiRP). *P* values in (d) and (e) are calculated by the t-test with Fischer transformation on Pearson’s *R.* jTiRP = TiRP (Treg-intrinsic regulatory potential) of the J-region of the TCR (IMGT positions 113-118) mTiRP = TiRP (Treg-intrinsic regulatory potential) of the middle region of the TCR (IMGT positions 105-112) vTiRP = TiRP (Treg-intrinsic regulatory potential) of the V-region of the TCR (IMGT positions 1-104)

PCA delineated two axes of TCR-driven cell states: antigen-experienced (Treg and memory Tconv) versus naïve (PC1), and regulatory versus conventional (PC2) (**Figure 7a**). The axis dividing antigen-experienced from inexperienced samples (PC1) was most reliant on *TRBV* gene frequencies, while the axis dividing regulatory versus conventional samples (PC2) was most reliant on mean percent composition of amino acids in CDR3βmr and the CDR3βmr-adjacent residue p113 (**Figure 7b-c**). Since TiRP is a weighted sum of TCR features from the V-, J- and middle regions, the score can be divided into three score components corresponding to these three regions. TiRP scoring by TCR region revealed that V-region-specific TiRP (vTiRP) and CDR3βmr-specific TiRP (mTiRP) indeed captured PC1 and PC2, respectively (**Figure 7d-e,** vTiRP – PC1 *R =* −0.86, *P =* 1.5 × 10^−20^, mTiRP – PC2 *R =* 0.85, *P =* 2.6 × 10^−20^).

We next investigated possible biological drivers for vTiRP and mTiRP. The biological structure of the pMHC-TCR complex suggests that different regions of the TCR may promote Treg fate via particular affinities: MHC II mostly contacts the V-region of the TCR, while the self-peptide is in closest contact with CDR3βmr^14,26,27^ (**Figure 1a**). Thus, we hypothesized that vTiRP was driven by affinity to human MHC II, while mTiRP was driven by affinity to self-peptides. To test this idea, we examined TiRP in two complementary datasets: 1) murine Treg TCRs^28^, which recognize self-peptide but not human MHC, and 2) human memory Tconv TCRs^12^, which recognize human MHC but not self-peptide (**Figure 8a**, **Table 1**).

**Figure 8.**
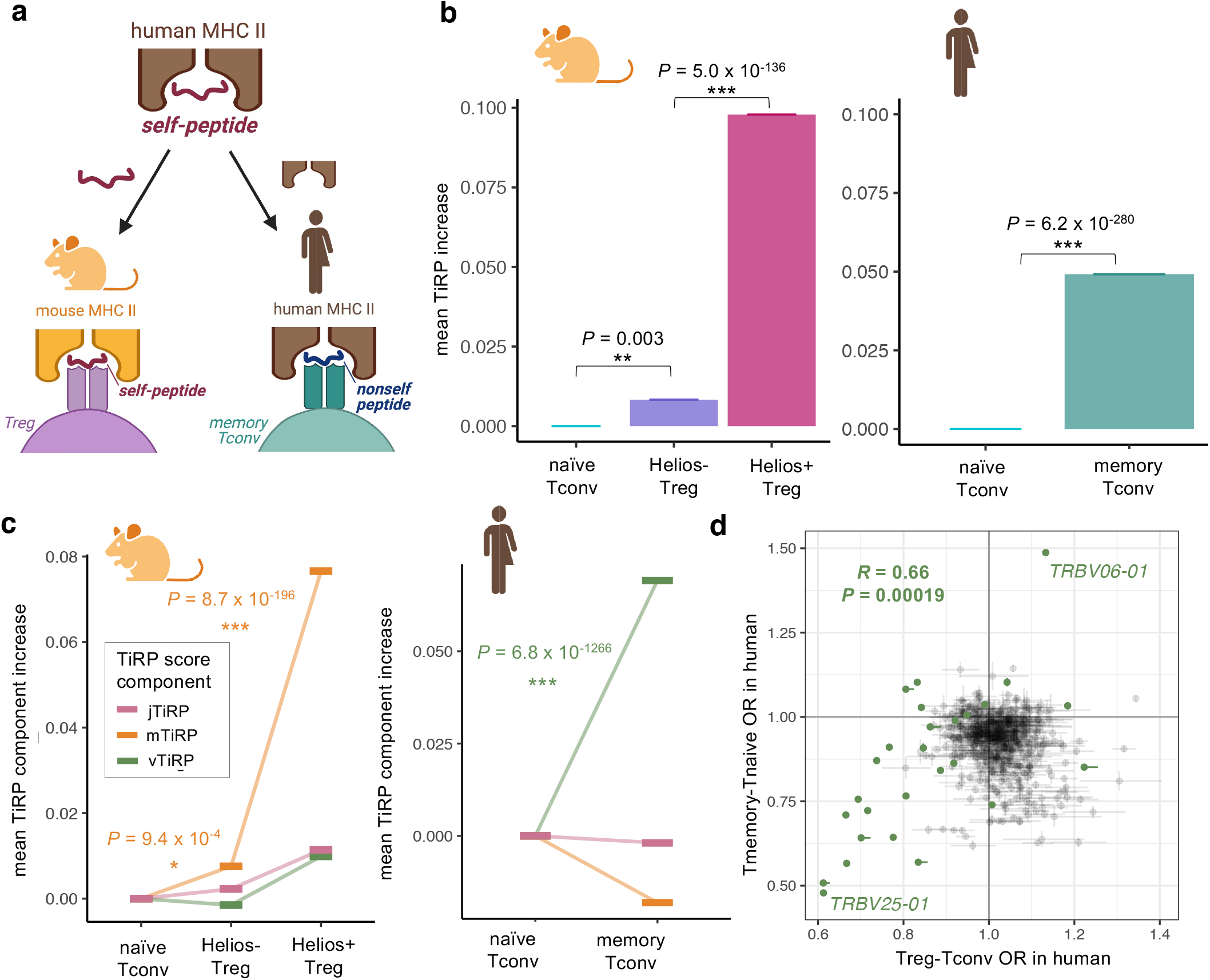
Isolating the drivers of TiRP. **(a)** We investigated the drivers of TiRP by separately examining the two elements of the human Treg TCR ligand: the self-peptide and the human MHC II molecule. To do so, we scored 1) murine Treg TCRs, which share an affinity to mammalian self-peptides but not to human MHC II molecules, and 2) human memory Tconvs TCRs, which share an affinity to human MHC II molecules but not to self-peptides. **(b)** Left: mean increase in TiRP score of *Helios*-sorted Tregs compared to naïve Tconvs in *Helios*-GFP *Foxp3*-RFP reporter mice. Right: mean increase in TiRP score of memory Tconvs compared to naïve Tconvs from held-out donors of the replication dataset. **(c)** Left: TiRP score increases in *Helios*-sorted murine Tregs broken down into TiRP score components by TCR region. Right: TiRP score increase in human memory Tconvs broken down into TiRP score components by TCR region. **(d)** Correspondence between TCR feature odds ratios for Treg-Tconv odds (x-axis, meta-analytic odds between discovery and replication cohort), and memory-naïve odds (y axis, replication cohort only) with their 95% confidence intervals. *TRBV* genes are highlighted in green. Pearson’s *R* is calculated with respect to *TRBV* gene odds ratios only. *P* values in (b)-(c) are calculated by LRT between mixed effects models (Methods); *P* value in (d) is calculated by the t-test with Fischer transformation on Pearson’s *R*. jTiRP = TiRP (Treg-intrinsic regulatory potential) of the J-region of the TCR (IMGT positions 113-118) mTiRP = TiRP (Treg-intrinsic regulatory potential) of the middle region of the TCR (IMGT positions 105-112) vTiRP = TiRP (Treg-intrinsic regulatory potential) of the V-region of the TCR (IMGT positions 1-104)

Applying our scoring system to murine data revealed that human TiRP was significantly elevated in murine Tregs compared to Tconvs (**Figure 8b**, left). Because the self-peptide is the only consistent element of the Treg ligand between these species, the best explanation for such cross-species Treg TCR similarity is affinity to thymic self-peptides. Indeed, TiRP was dramatically elevated in murine Tregs that expressed *Helios*, a marker of thymic Treg fate acquisition (**Figure 8b**, left). Consistent with our TCR region hypothesis, the TiRP component with the greatest increase between murine Tconvs and Tregs was mTiRP (**Figure 8c**, left). CDR3βmr amino acid percentage effect sizes replicated strongly between murine and human data (**Extended Data Figure 8c**) while other TCR features did not (Pearson’s *R =* 0.85, *P =* 0.00013 for CDR3βmr amino acid percentages, **Extended Data Figure 8d, Supplementary Table 10, Methods**). These results strongly suggest that CDR3βmr features such as hydrophobicity promote Treg fate via affinity to self-peptide.

To understand the role of human MHC, we compared TiRP in naïve and memory Tconv TCRs^12^, which do not strongly recognize self-peptides^6^ (**Figure 8a**, **Table 1**, **Methods**). TiRP was significantly elevated in human memory Tconvs compared to human naïve Tconvs (**Figure 8b**, right), indicating that affinity to human MHC II also contributes to TiRP. Consistent with the hypothesis of V-region-based affinity to human MHC II molecules, vTiRP was the only TiRP component to increase in human memory Tconvs (**Figure 8c**, right). As expected, large-effect size TCR features between memory Tconvs and naïve Tconvs were predominantly *TRBV* genes (**Figure 8d**, **Extended Data Figure 8e**), and the extent of each gene’s enrichment in memory Tconvs correlated with the extent of its enrichment in Tregs (**Figure 8d,** Pearson’s *R =* 0.702, *P =* 4.5 × 10^−5^ for *TRBV* genes). These effects further replicated in an entirely independent cohort of sorted memory and naïve T cells from 5 healthy donors^29^ (**Table 1, Extended Data Figure 8f, Supplementary Table 11**). Thus, as structural interactions in the pMHC-TCR complex would suggest, V-region features modulate affinity to MHC, thereby shaping the T cell’s general disposition for activation.

## DISCUSSION

The majority of regulatory T cell research to date has focused on TCR-extrinsic determinants of T cell phenotype, such as the effect of costimulatory receptors, antigenic peptides, and cytokines^30^. Though each of these elements certainly play an essential role in T cell fate, the contribution of the TCR sequence itself has not yet been comprehensively investigated. TCR-intrinsic factors are relevant to nearly all immunological contexts, including the engineering of TCRs for immune therapies. The presence of TCR sequence features that generally promote TCR signaling in engineered TCRs would similarly make T cell activation more likely, regardless of host factors. Because TCR sequence arises from a random process prior to T cell fate determination, TCR features must be causal for T cell fate.

In this work, we leveraged the affinity-based partition of the repertoire into Tregs and Tconvs to uncover determinants of TCR avidity towards self-antigens. We identified robust effects of TCR sequence that are predictive of T cell fate across six independent cohorts, encompassing diverse genetic, clinical and tissue contexts. Our results were robust to different sequencing protocols used across these studies, suggesting that our findings are robust to technical factors. Using the comprehensive set of predictive TCR sequence features, we developed and validated a score capturing the TCR-intrinsic regulatory potential (TiRP) in the V-region, CDR3 middle region, and J-region. Excitingly, this score helped to explain the tendency for expanded tumor-infiltrating T cell clones to adopt a regulatory phenotype.

It is important to recognize several limitations to our approach. First, the amount variance in T cell state explained by the TCR is significant but modest considering the full diversity of the repertoire. For any given TCR, specific antigenic contacts and costimulatory signals are likely the major determinants of T cell phenotype. Our results show, however, that TCR features such as hydrophobicity consistently predispose the T cell to adopt a regulatory phenotype. Second, our analyses focused on the β chain of the TCR. The β chain is more variable than the α chain and is largely considered to mediate antigen specificity. However, the α chain may also play a role in determining T cell phenotype, which remains to be explored. Furthermore, sequence features of the β and α chains may act synergistically and the combination of data from both may be more highly predictive than the β chain alone. Lastly, though our questions focused on thymic biology, the TiRP scoring system is based on TCRs sampled from extrathymic tissues. Since the majority of circulating Tregs are thought to be thymically-derived^28,31,32^, we suspected that extrathymic Treg-Tconv phenotypes would mostly reflect thymic T cell fate determination. Indeed, our assessment of TiRP scores in Helios-GFP Foxp3-RFP mice argue that TiRP is specifically elevated in thymically-derived Tregs.

The broadest takeaway from our work is the hydrophobic bias of Treg TCRs, present at each of the entropic positions of CDR3β. This observation extends previous work^15,33^ regarding p109 and p110 of Treg TCRs, and demonstrates that the hydrophobic bias is in fact not position-specific. By the hydrophobic effect, hydrophobic amino acids aggregate into high affinity hotspots, which may afford a degenerate “stickiness” to hydrophobic TCRs. By this explanation, CDR3β hydrophobicity may increase the likelihood of Treg fate not by extreme affinity to particular self-peptides, but rather by minimal affinity to a larger pool of cognate antigens. Thus, via the flexibility of a hydrophobic TCR, the Treg may generalize from the self-peptide encountered in the thymus to a larger pool of protected self-antigens. Though focused on CD8^+^ T cell biology, the enhanced immunogenicity of hydrophobic CD8^+^ T cell epitopes^34^ may reflect a similar underlying concept: the best attractor to a large pool of diverse cognate ligands (in this case, the CD8+ repertoire) is the broadly-interacting feature of hydrophobicity.

Importantly, however, CDR3β hydrophobicity is not the full picture. *TRBV* gene usage explained nearly as much variance in T cell fate, and *TRBV* gene effects were not related to hydrophobicity. Our work suggested instead that the isoelectric point of the CDR1β p37 encoded by the *TRBV* gene shapes affinity to conserved sites of MHC II^14^. We observed *TRBV* gene biases pertained to memory Tconvs as well, indicating that some *TRBV* genes provide enhanced, antigen-invariant affinity to MHC II that predisposes the TCR for activation.

These phenomena offer a new lens on the T cell immune response: though each TCR tends to recognize a specific cognate antigen, all TCRs are subject to common processes that shape T cell activation. Due to these common processes, not all TCRs are created equal—those with a higher baseline for general reactivity may require a less “perfect” cognate antigen for activation. Existing tools provide rough annotations for “TCR strength,” but these are based on frequently interacting residues in general protein structures^35^. TiRP sharpens our understanding of high affinity amino acids in the context of the pMHC-TCR complex, providing a crucial functional annotation for the T cell receptor.

## ONLINE METHODS

### Bulk sequencing data

We downloaded the discovery cohort^11^, replication cohort^12^, the murine cohort^28^ and memory cohort^29^ sequencing data from the Adaptive Biotechnologies immuneACCESS site (https://clients.adaptivebiotech.com/immuneaccess). For all data, we defined CDR3 amino acid sequences with stop codons or frameshifts to be non-productive amino acid sequences. We restricted all analyses to CDR3 sequences of a length within 12 and 17 amino acids, representing 91.8% of observations in the discovery cohort. We aligned CDR3 amino acids to positions defined by IMGT^37^, wherein sequences less than 15 amino acids have mid-region gaps and sequences longer than 15 amino acids have extra mid-region positions. We examined only one copy of each CDR3β sequence within each individual. Unless explicitly noted, we excluded CDR3β reads that were observed in both the Treg and Tconv sample of any individual (0.63% of observations in the discovery cohort and 1.9% of observations in the replication cohort). We excluded two individuals in the discovery cohort (donor IDs = 6174 and 6282) and 12 individuals in the replication cohort (donors IDs = HD9, HD10, HD11, HD12, HD13, HD14, T1D1, T1D9, T1D10, T1D11, T1D12, T1D14) because these donors did not have both Tregs and Tconvs available.

### Single cell sequencing data

We downloaded scRNAseq tumor microenvironment data from the GEO through accession numbers GSE114727, GSE114724, and GSE123814. For quality control, we included only cells for which 1) more than 1000 genes were expressed 2) less than 25% of detected UMIs were of mitochondrial origin and 3) exactly one productive TCR beta chain was detected. We followed the quality control process of the original authors for the multimodal memory T cell dataset^25^, which is available for download from the GEO through accession number GSE158769.

## STATISTICAL ANALYSES

All mixed effects models were fit with R package lme4. All model comparisons were computed with R package anova. All significance tests on Pearson’s *r* were t-tests with the Fischer transformation. All analyses were done with R version 3.6.1.

### Holding out observations for calibration and testing

To leverage both the discovery (Seay et al. 2016) and replication (Gomez-Tourino et al. 2017) cohorts in the development of TiRP, we used approximately 70% of the TCR clones from each cohort for training, 10% for calibration, and 20% for testing. To preserve the novelty of held-out data, we kept all TCR clone observations from the same individual together in this process, holding out entire repertoire samples. In the discovery cohort, we held out two individuals for TiRP calibration (donor IDs = 6279, 6196, accounting for 8.4% of TCR clones in the discovery cohort) and four individuals (donor IDs = 6161, 6193, 6207, 6287, accounting for 20.3% of clones in the discovery cohort) for TiRP testing. In the replication cohort, we held out one individual for TiRP calibration (T1D3) and three individuals (HD1, HD2, T1D6) for validation. TCR sequence feature effect sizes were estimated in a separate mixed effects model for each cohort for each independent region of the TCR.

### Mutual information structure of the CDR3β sequence

We calculated the Shannon entropy^38^ of each CDR3 position and the mutual information^39^ between all pairs of CDR3 positions with the R package DescTools. To normalize the mutual information, we divided the mutual information by the entropy at each position, and then took the harmonic mean of these two ‘coefficients of constraint’^40–42^.

### Selection of random effects and model comparisons

In the discovery cohort^11^, T cells were sampled from four tissues: peripheral blood (PBMC), spleen, pancreatic lymph node (pLN), and inguinal/irrelevant lymph node (iLN). We reasoned that there were three sensible ways to model tissue as a source of CDR3β variation: (1) as a fixed effect:

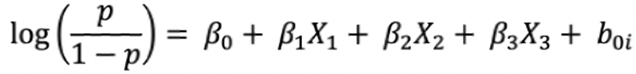

where *p* is the probability that the CD4+ sorted CDR3β sequence belongs to a Treg, *β_0_* is an intercept, *X_1_* is an indicator variable set to 1 if the sequence is from a PBMC sample, *X_2_* is an indicator variable for spleen origin, *X_3_* is an indicator variable for iLN origin (pLN as reference), and *b*_1*i*_ is a modification to the intercept fit to each individual *i*, normally and identically distributed (NID) with mean 0 and variance σ_0_^2^.

(2) as a random intercept effect independent from the random intercept effect per individual, wherein matched tissues across donors have the same (zero-centered) intercept effect:

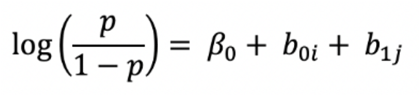

where *b*_1*j*_ is a modification to the intercept fit to each tissue *j*, NID with mean 0 and variance σ_1_^2^, and all other variables maintain previous definitions

and/or (3) as a nested random intercept effect, wherein each tissue-donor pair is modeled as a unique batch of correlated observations within the individual-level and tissue-level variances:

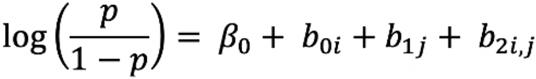

where *b*_2*i,j*_ is a modification to the intercept fit to each individual *i* - tissue *j* pair, NID with mean 0 and variance σ_2_^2^, and all other variables maintain previous definitions. For stable numerical results, we included the marginal random effects for donor and tissue in this nested random intercept model.

To determine which of these models was most appropriate, we calculated the pseudo *R*^2^ by the conventional McFadden^43^ approach (range 0-1). All measures of variance explained in this study were computed with this approach. For this analysis, we compared models 1-3 to a baseline model that fit the log odds of Treg status only to a random intercept for each individual:

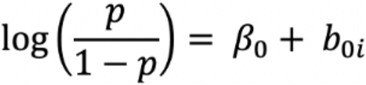

These model comparisons revealed that tissue explained 1.90% of variance as a fixed effect and 1.15% of variance as a random effect (*P =* 1.15 × 10^−11211^ fixed and *P =* 4.68 × 10^−10229^ random, LRT). On the other hand, tissue as a random effect nested within individual explained 6.27% of variance (*P =* 1.32 × 10^−55291^, LRT). We therefore concluded that nesting a random tissue effect within the donor random effect was the most appropriate model for the batch structure of these data, and proceeded with three random intercepts for each mixed effects model: the nested donor-tissue effect, the marginal donor effect, and the marginal tissue effect.

### CDR3βmr mixed effects logistic regression

For each amino acid, we calculated the percentage of CDR3βmr positions occupied by this residue; a percentage of 0 means that the residue is missing for a given TCR, while a percentage of 100 means that the residue is present at every CDR3βmr position. We scaled this percentage to have a mean of 0 and variance of 1, and tested the scaled percentage in a separate mixed effects logistic regression for each amino acid with random intercepts as described above. We controlled for CDR3β sequence length by including it as a categorical covariate, reasoning that conformational differences in the HLA-TCR complex may not scale linearly with additional residues. To collect the relevant amino acid proportions, we did a forward search where we iteratively added to the mixed effects model the amino acid proportion that provided the greatest improvement in model fit. On the first round, the percentage of CDR3βmr positions occupied by Glutamic acid (E) in each TCR explained the most variance, with a 9.7% fall in odds of Treg fate per additional Glu residue for CDR3βs of length 15 (pseudo R^2^ = 0.036%, likelihood ratio test (LRT) *P =* 8.37 × 10^−196^, OR = 0.954, 95% CI = 0.951 – 0.957). Conditioning on this feature revealed that the next amino acid with the greatest independent effect was Aspartic acid (D) (pseudo R^2^ = 0.042%, LRT *P =* 1.01 × 10^−225^, OR = 0.95, 95% CI = 0.947 – 0.953). We repeated this process until the remaining amino acid percentages no longer passed the Bonferroni-corrected significance threshold (p=0.05/20 amino acids) (**Figure 3b**, middle). We confirmed that this threshold kept the type I error rate below 0.05 by repeating this analysis 1000 times, with Tconv and Treg labels for each TCR randomly shuffled within the data for each donor on each run.

### Position-specific mixed effects logistic regressions

To parse the *TRBV-*encoded region, we asked if the 5’ flanking CDR3β residues could be represented by a handful of motifs. Indeed, the 8 p104-p106 sequences (“Vmotifs”) present in each donor with frequency > 0.001 in every donor accounted for 96.2% of TCRs. We labeled the remaining 3.8% of TCRs with a Vmotif of “other” and assessed the association between Vmotif and T cell fate with a mixed effects model including p107 as a fixed covariate. We found that the Vmotif indeed explained significant variance in Treg fate (pseudo R^2^ = 0.02%, *P =* 3.57 × 10^−93^, LRT). However, because the Vmotif strongly corresponds to *TRBV* gene usage (**Extended Data Figure 1**), we next evaluated whether Vmotif effects are in fact mediated by their corresponding *TRBV* genes. Indeed, adding *TRBV* gene identity to the mixed effects model as a fixed covariate abrogated the significance of the Vmotif term, but not the p107 term. For this reason, we concluded that the *TRBV*-encoded region was best modeled by joint estimation of *TRBV* gene and p107 residue effect sizes, with donor-individualized *TRBV* gene thymic selection rate as a fixed covariate (**Supplementary Note**).

Similarly, to parse the *TRBJ-*encoded region, we asked if the 3’ flanking CDR3β residues could be represented by a handful of motifs. Indeed, the 42 p114-p118 sequences (“Jmotifs”) present in each donor with frequency > 0.001 in every donor accounted for 91.5% of TCRs. In analogous model comparisons, donor-individualized *TRBJ* gene selection rates did not explain a significant amount of variance in Treg fate, but the Jmotif, *TRBJ* gene, and the p113 residue each did. In contrast to the *TRBVregion*, here it appeared that the motif mediated the effect of the gene, with the Jmotif explaining slightly more variance than the *TRBJ* gene (**Figure 3b**, right). Thus, we concluded that the *TRBJ*-encoded region was best modeled by joint estimation of Jmotif and p113 residue effect sizes.

To protect against numerically unstable estimates, we report only the effect sizes of TCR features with a frequency greater than 0.005 in the training data for both the discovery and replication cohorts.

### Estimating the effects of physicochemical features

To estimate the effects of physicochemical features, we represented each CDRβ loop residue as a vector of length 3, corresponding to the amino acid’s hydrophobicity, isoelectric point, and volume. For consistency with the closely related work by Stadinksi et al.^15^, we used the whole-residue interfacial hydrophobicity scale^44^. Isoelectric point values were obtained from the CRC Handbook of Chemistry and Physics^45^ and volume measurements were obtained from Zamayatnin 1972^46^. Each value was scaled to have a mean 0 and variance 1 in the discovery cohort training data.

To localize the importance of these physicochemical features within the TCR, we represented each residue belonging to a CDRβ loop as a vector of length 3 corresponding to the amino acid’s hydrophobicity, isoelectric point, and volume, and modeled Treg fate as an outcome of these features using multiple logistic regression. We followed IMGT positioning, wherein the human CDR1β loop consists of positions 27à 38; while the human CDR2β loop consists of positions 56 à 65. We used only TCR reads with a resolved *TRBV* gene (78.5% of observations), and imputed CDR loop amino acids based on *TRBV* gene identity using IMGT^37^. To enable TCR alignment, we discarded 3.6% of observations with a resolved *TRBV* gene for which there were not exactly 5 CDR1β amino acids and 6 CDR2β amino acids, or for which CDR1-2 amino acids were not available via IMGT.

To handle the densely correlated TCR features within these loops, we applied a ridge penalty to the logistic regression using R package “glmnet.” This coefficient penalization obviated the need for random effects, and so we included batch (donor and tissue source of the TCR) as a fixed and penalized covariate. As in the *TRBV* gene analysis, we used VGSR as a covariate to partial out genetic variation in *TRBV*-MHC affinity (**Supplementary Note**). All predictors were scaled to a have mean 0 and variance 1. We did not assume that position-wise physicochemical effects would translate across different CDR3β lengths, and so fit a separate logistic regression for each length. For each regression, we tuned the lambda shrinkage penalty by testing the 100 values generated by the glmnet package and selecting the one that gave the minimum mean cross-validated error across 10 folds of the training data in the discovery cohort.

In a separate analysis isolated to the CDR3βmr, we fit a separate mixed effects logistic regression for each length-position combination in the discovery cohort training data (**Extended Data Figure 5b**). We included all three physicochemical features as fixed covariates for each position, and modeled donor and tissue sources as random effects as described above. Each physicochemical feature was scaled to have a mean 0 and variance 1 for each length-position combination.

For the **Figure 4c** visualization, we included only TCRs with a CDR3β length of 15 amino acids in the discovery cohort training data, and fit a separate mixed effects logistic regression for each position. Each regression included random intercepts as described above and one fixed covariate corresponding to the amino acid identity at the given position. We cast the most common amino acid as the reference: Leucine for position 108, and Glycine for all other positions.

### Developing the TiRP scoring system

We defined TiRP as the sum of the TCR sequence features present in a given TCR, reasoning that the effects of TCR features were additive provided that they were fit jointly or derived from independent regions of the TCR. To reach a consensus effect size for each TCR feature across the two cohorts, we used inverse-variance weighted meta-analysis (meta-analytic effect size for feature *X =* average of the discovery cohort and replication cohort effect sizes for feature *X*, weighted by their respective standard errors). Due to the inconsistent effect size directions for %V in the CDR3βmr (**Figure 5a**, **Extended Data Figure 3b**), we included only 14 amino acid percent covariates in our final CDR3βmr models (**Supplementary Table 1, Supplementary Table 7**). To exclude potentially unreliable effect size estimates from the score computation, we calibrated a meta-*P* value significance threshold above which TCR features were excluded from the score. For this, we used a single mixed effects logistic regression for each threshold over a range of thresholds on the pooled discovery and replication TCRs held out for calibration (discovery cohort: 6279, 6196, replication cohort: T1D3). Each mixed effects logistic regression estimated the fixed effect of TiRP on T cell fate, with random intercepts for donor source, tissue source, and each donor-tissue source pair (see “selection of random effects and model comparisons”). We found that no threshold explained significantly greater variance than the Bonferroni-corrected threshold, 0.05/612 TCR features, resulting in 25 *TRBV* genes, 23 Jmotifs, 4 CDR3β lengths, 14 CDR3βmr amino acid percentages, and 143 position-specific features relevant to TiRP computation (**Supplementary Table 7**).

### Testing TiRP in held-out donors from bulk sequencing cohorts

To test TiRP in bulk sequencing data, we scored each unique productive TCR in donors held out from both TiRP training and calibration (discovery cohort donors 6161, 6193, 6207 and 6287, and replication cohort donors HD1, HD2, and T1D6). We then tested the association between TiRP and T cell state by comparing the additional variance explained by a mixed effects logistic regression model including TiRP as a fixed covariate to a baseline model containing only donor ID, tissue source, and donor-tissue interaction as random intercepts (likelihood ratio test). We conducted the same process for nonproductive TCRs in held-out donors, and restricted this analysis to the discovery cohort, in which TCR gDNA was sequenced and therefore out-of-frame reads were available (**Table 1**). To ascertain the difference between high-scoring and low-scoring TCRs in these held-out data, we collected the top and bottom decile of TCRs per donor, and compared the ratio of Tregs to Tconvs between the group of all top decile TCRs and the group of all bottom decile TCRs.

### Validating TiRP in single-cell data of tumor microenvironment

In single-cell data analyses, TCR clones were defined by a barcode consisting of their donor ID and CDR3β DNA sequence. As in bulk sequencing analyses, CDR3β chains with a length shorter than 12 amino acids or longer than 17 amino acids were discarded. Only cells with exactly one productive CDR3β detected were included in analyses.

We computed the TiRP score for each clone based on its CDR3β amino acid sequence and *TRBV* gene. So that TiRP scores would be comparable, percent amino acid values were scaled by the mean and standard deviations of the TCRs held out for testing from the discovery and replication cohorts (transformation provided in Supplementary Table 3). *TRBV* gene usage was determined by MixCR alignments for the Azizi et al. cohort and RNA expression in the Yost et al. cohort. To determine *TRBV* gene usage based on RNA expression in the Yost et al. cohort, read counts were log-normalized per cell and then scaled so that each *TRBV* gene had mean 0 and variance 1 within cells that had non-zero read counts for the given gene. Each cell was then assigned the *TRBV* gene with the highest normalized and scaled expression. Cells without any *TRBV* gene expression detected were given a *TRBV* gene value “unresolved.”

To validate the TiRP score in these data, we tested the association between TiRP score and regulatory or conventional cell phenotype. For the Yost et al. cohort, cell phenotypes based on the original authors’ clustering were available. We labeled all cells in the ‘Tregs” and “Treg” cluster as regulatory and all cells in the “Tfh”, “Th17”, “CD4_T_cells”, and “Naïve” to be CD4^+^ conventional. Because original authors’ cell phenotype labels were not available for the Azizi et al. cohort, we applied a standard scRNAseq pipeline to infer cell phenotypes: we excluded all cells with read counts from 1000 genes or less or at least 25% of read counts from mitochondrial genes and then used Seurat^47^ with default parameters to 1) normalize the read counts per cell, 2) take the variance-stabilizing transform 3) scale and center gene expression, 4) compute the first 20 principal components based on the 500 most variable genes, 5) harmonize the principal component embeddings by sample (donor_batch ID) with R package “harmony” using default parameters, 6) construct a shared-nearest-neighbor (SNN) graph based on these harmonized embeddings with k=30, 7) conduct Louvain clustering on the SNN graph with resolution 0.8, and 8) run uniform maniform approximation and projection on the first 10 harmonized PCs.

### Creating a CD4+ memory T cell single cell reference

To construct a reference of cellular phenotypes for CD4+ memory T cells, we used a published dataset^25^ of scRNAseq and CITE-seq for 500,000 memory T cells from 259 donors (**Table 1**). From these quality-controlled data, we used CITE-seq values to select 430,270 CD4+ cells (normalized CD4 > 1.5 and normalized CD8 <1, consistent with the original authors’ procedure). We followed the method developed by Nathan et al. to cluster the cells based on integrated mRNA and protein expression. First, we used Seurat^47^ to normalize the read counts per cell, take the variance-stabilizing transform and then scale gene expression to have a mean 0 and variance 1. We selected the union of the 1500 most variable genes (by mRNA expression) in each donor, resulting in 4707 variable genes.

To integrate surface protein information, we used CCA. First, we resolved the coefficients that maximized the correlation between linear combinations of the 4707 genes and the 31 manually-curated surface proteins^25^ in the CITE-seq panel (“cc” function from R package “CCA”). We then projected the cells into the 31 canonical dimensions in mRNA space, and used Harmony with default parameters to harmonize the embeddings of these canonical dimensions by donor. For visualization, we used the R package uwot to conduct UMAP on the first 10 canonical dimensions using the cosine metric, a local neighborhood size of 30, and a minimum distance of 0.3 between embeddings. To identify cell types, we constructed a SNN graph (k=10) from the harmonized embeddings of the first 10 canonical dimensions, and conducted Louvain clustering on the SNN graph with resolution 0.8, revealing one cluster (#6) with markedly elevated *FOXP3* and CD25 expression and reduced CD127 expression. We labeled cells belonging to this cluster as Tregs and manually annotated the phenotypes of the other clusters based on surface expression of the 31 manually-curated, immunologically relevant surface proteins as well as mRNA expression of *CCR7, IFNG, GZMK,* and *CTLA4* (**Extended Data Figure 6c-d**).

### Mapping tumor-infiltrating T cells with Symphony

Before ascertaining mixed clones in tumor-infiltrating cells, we standardized Treg and Tconv definitions between the two cohorts by projecting cells from both cohorts into the annotated low-dimensional space of the reference single cell dataset. To accomplish this projection and simultaneously harmonize the tumor-infiltrating cells by cohort, donor and sample, we utilized Symphony^24^. Because the reference dataset consisted of only memory T cells and our hypothesis focused on expanded clones, we mapped only the tumor-infiltrating cells for which their paired CDR3β DNA sequence was detected on more than one cell within their patient sample (56.1% of cells in the Azizi et al. cohort, 60.6% of cells in the Yost et al. BCC cohort, and 73.7% of cells in the Yost et al. SCC cohort). For each cohort separately, we used Symphony to map the query cells into the harmonized reference canonical variate embedding space while integrating over unwanted sources of technical variation tagged by donor and sample in the query. We used the resultant canonical variate embeddings to 1) impute cluster membership for query cells via k-nearest-neighbors in the reference cohort (R package “knn”, k=5), and 2) project the query cells into the reference UMAP embedding. To visualize TiRP trends, we colored each cell by the average TiRP of its 100 nearest query neighbors in the 31 canonical dimensions (**Figure 6c**).

### Mixed clone analysis with bulk sequencing data

We conducted our mixed clone analysis with bulk sequencing data in the donors from the discovery and replication cohort that were held out from the estimation of TCR feature effect sizes and TiRP score calibration (**Figure 1b**, **Table 1**). Clones were defined by the “barcode” consisting of their CDR3β nucleotide sequence, *TRBV* gene ID, and donor ID. Because clonal expansion is a prerequisite to mixed clone status, we compared mixed clone TiRP scores to those of expanded Tconv and Treg clones. For the discovery cohort, *TRB* chains were sequenced from gDNA, and so clonal expansion could be derived from the number of “templates” for each clone (number of biological molecules prior to PCR amplification, inferred by immunoSEQ via internal bias control). Because *TRB* chains were sequenced from cDNA in the replication cohort, we cannot know whether identical reads within the same sample represent *TRB* transcripts from one or multiple cells. However, we can deduce that identical reads across multiple FACS-sorted samples from the same individual arose from multiple cells and therefore an expanded clone. Therefore, for the replication cohort, we collected a sample of the expanded clones from each donor by aggregating all CDR3β nucleotide sequences that arose in multiple FACS-sorted samples from the same individual (Treg, naïve Tconv, central memory Tconv, and stem-cell like memory Tconv). Because there was only one Treg sorted sample for each individual, we could only detect pure Tconv or mixed clones in the replication cohort. We tested the effect of TiRP score on clone phenotype with mixed effects models as designed in the single-cell analyses.

### Mixed clone analysis with single cell data

To detect mixed clones in single cell data, we aggregated cells into clones based on matching clonal “barcodes:” patient ID, *TRB* DNA sequence, *TRBV* gene, and TRA amino acid sequence. To protect against contamination by doublets (droplets encapsulating two cells rather than one), we excluded cells with more than one unique TRB chain detected. Since the expression of multiple TRA chains, however, is a common biological phenomenon^48^, we did not exclude multi-TRA chain cells. To assign a clonal barcode TRA for these cells, we selected the TRA sequence that was most often expressed by cells with a matching *TRB* DNA sequence in the given patient.

To model the effect of TiRP score on clone phenotype (Tconv, Treg, or mixed), we used mixed effects logistic regression with random intercept for the clone’s source patient and the clone’s source cohort (BRCA, SCC, or BCC). Since clonal expansion is a prerequisite to mixed clone status, only clones of size > 1 were included. We used the LRT to compare the model including TiRP to a baseline model containing only the random covariates. We conducted this process twice: first to compare mixed clones to purely Tconv clones, and second to compare mixed clones to purely Treg clones.

We then quantified the clone phenotype by taking the natural log transform of the within-clone Treg/Tconv ratio, with one “hallucinated” Treg and one “hallucinated” Tconv per clone to protect against numerically unstable estimates. We tested the effect of TiRP score on this quantitative clone phenotype using mixed effects linear regression with random intercepts as described above, and found a 0.065 increase in ln(Treg/Tconv ratio) per standard deviation increase in TiRP score (**Figure 6h**, *P =* 1.6 × 10^−4^, LRT).

To check that *FOXP3* expression was significantly different between Tregs and Tconvs within mixed clones, we conducted a Student’s paired t-test and confirmed that this was indeed true (**Extended Data Figure 7e**).

### Analysis of murine TCRs

T cell clones were defined by the barcode consisting of CDR3β amino acid sequence, *TRBV* gene identity, and donor ID. Due to ambiguity, clones observed in both Treg and Tconv samples from the same donor or in both the Helios+ and Helios- Treg samples from the same donor were excluded from the following analyses. Clones with member cells in both the naïve Tconv and memory Tconv samples from the same donor were labeled with the memory Tconv phenotype.

To compute the *TRBV* gene component of the TiRP score in murine data, we assigned each murine *TRBV* gene the TiRP coefficient of its human homolog according to human-mouse *TRBV* correspondences listed in IMGT^37^. Murine and human *TRBV* genes were aligned for comparison in **Extended Data Figure 8d** by this same correspondence scheme. Murine *TRBV* genes with multiple human *TRBV* gene homologs were assigned the average of their human homolog coefficients. Because the reference *TRBV* gene in human data, *TRBV05-01*, does not have a murine homolog, comparing *TRBV* gene effect sizes in mouse and human required a change to a common reference. We encoded *TRBV19-01* as the reference for murine mixed effects logistic regression models, and translated human *TRBV* gene effect sizes to those that would be obtained from *TRBV19-01* as the reference by subtracting the meta-analytic effect size for *TRBV19-01* from all *TRBV* gene effect sizes (including *TRBV05-01*, originally at 0).

### TCR feature Principal Components Analysis

To contextualize the amount of T cell phenotypic variation explained by TCR features identified in our work, we performed a principal components analysis on the matrix of samples by TCR feature means for the replication cohort, in which sorted samples for all T cell phenotypes of interest were available (**Table 1**, **Figure 7a**). For categorical TCR features such as *TRBV* gene or Jmotif, we one-hot-encoded the variable into a binary vector equal to the length of possible values, and took the mean of each of the positions. As this process rapidly expands the dimensionality of each sample, we summarized the TCR features in the CDR3βmr by percent composition of each amino acid only. We used the function “prcomp” from R package “stats” to conduct singular value decomposition of the centered and scaled matrix of samples by mean TCR features.

### Memory-Naïve TCR comparisons

T cell clones were defined by the barcode consisting of CDR3β amino acid sequence, *TRBV* gene identity, and donor ID. Due to ambiguity, clones observed in both Treg and Tconv samples from the same donor were excluded from the following analyses. Clones with member cells in both the naïve Tconv and memory Tconv samples from the same donor were labeled with the memory Tconv phenotype.

For the replication of Tconv Memory-Naïve TRBV effects in the Soto et al. cohort^29^, two additional steps were necessary to accommodate the deeper TCR sequencing within these individuals. First, only TCRs with a Cysteine at position 104 and Phenylalanine at position 118 were included. Though there does exist some minor physiologic variation at these conserved sites, such outlier sequences are not relevant to TiRP score computation. Second, though the donor source of each TCR was modeled as a random effect in other cohorts, we modeled it here as a fixed covariate, reducing computational burden and allowing the maximum likelihood estimation to converge.

## Acknowledgments

We thank Michael B. Brenner for helpful scientific conversations regarding this work.

K.A. Lagattuta and J.B. Kang are each supported by award number T32GM007753 from the National Institute of General Medical Sciences.

A. Nathan is supported by award number T32AR007530 from the National Institute of General Medical Sciences.

D.A. Rao is supported by NIH NIAMS K08 AR072791 and a Career Award for Medical Sciences from the Burroughs Wellcome Fund.

## Author contributions

K.A. Lagattuta, K. Ishigaki, and S. Raychaudhuri conceived the study. K.A. Lagattuta performed computational analyses. All authors contributed to data interpretation. K.A. Lagattuta, K. Ishigaki, and S. Raychaudhuri contributed to writing the manuscript. All authors reviewed the manuscript. K. Ishigaki and S. Raychaudhuri supervised the study.

## Code availability

Upon acceptance, scripts to reproduce analyses will be made available on GitHub.

