## Extended Data Figures for "Repertoire analyses reveal TCR sequence features that influence T cell fate"

### Extended Data Figure 1.

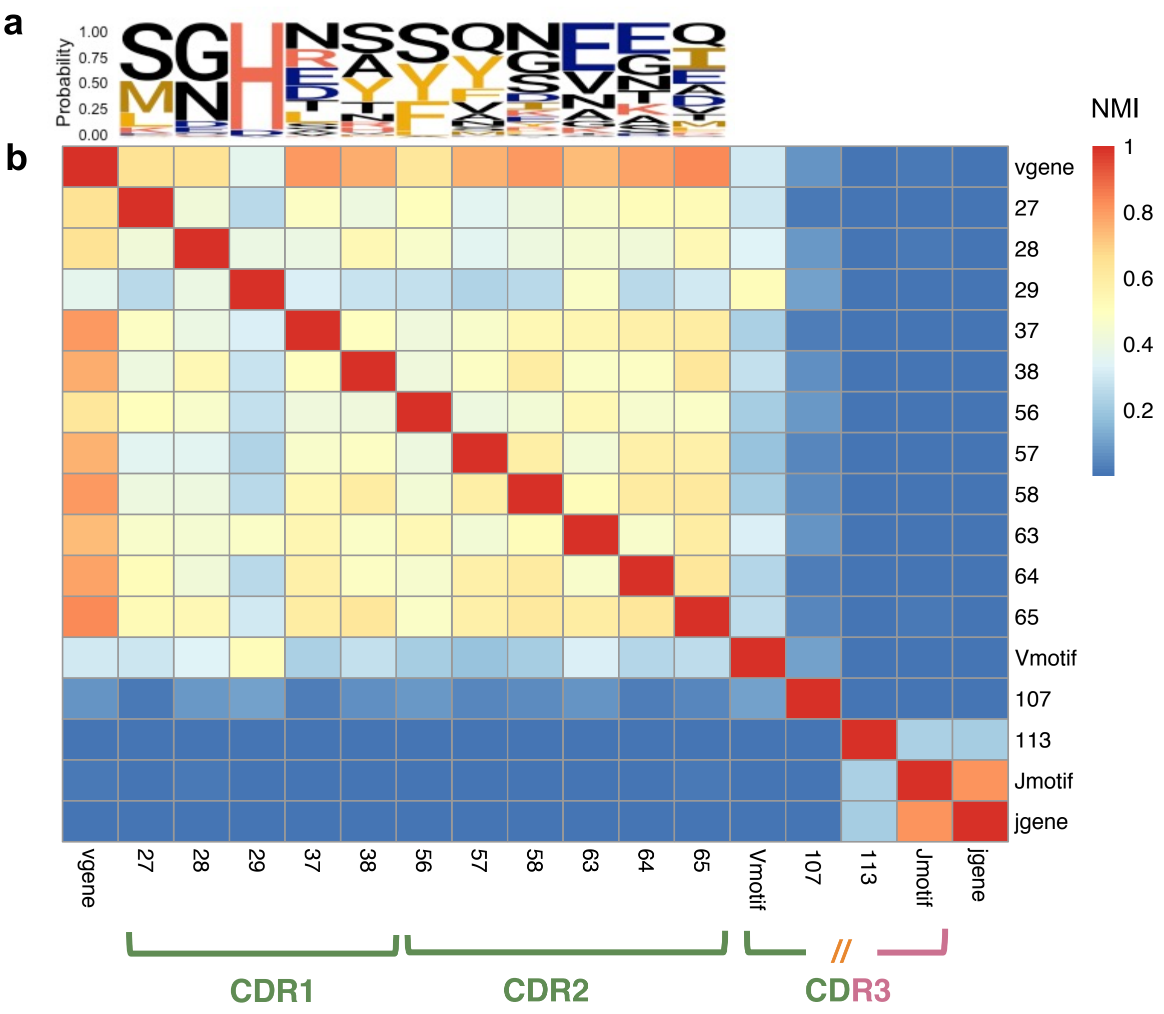


**Extended Data Figure 1. (a)** Probability of each amino acid in each CDR(1-2)β loop position, with **(b)** normalized mutual information between each pair of CDR(1-2)β residues, and *TRB(V/J)*-derived sequence features in the discovery cohort.

### Extended Data Figure 2.

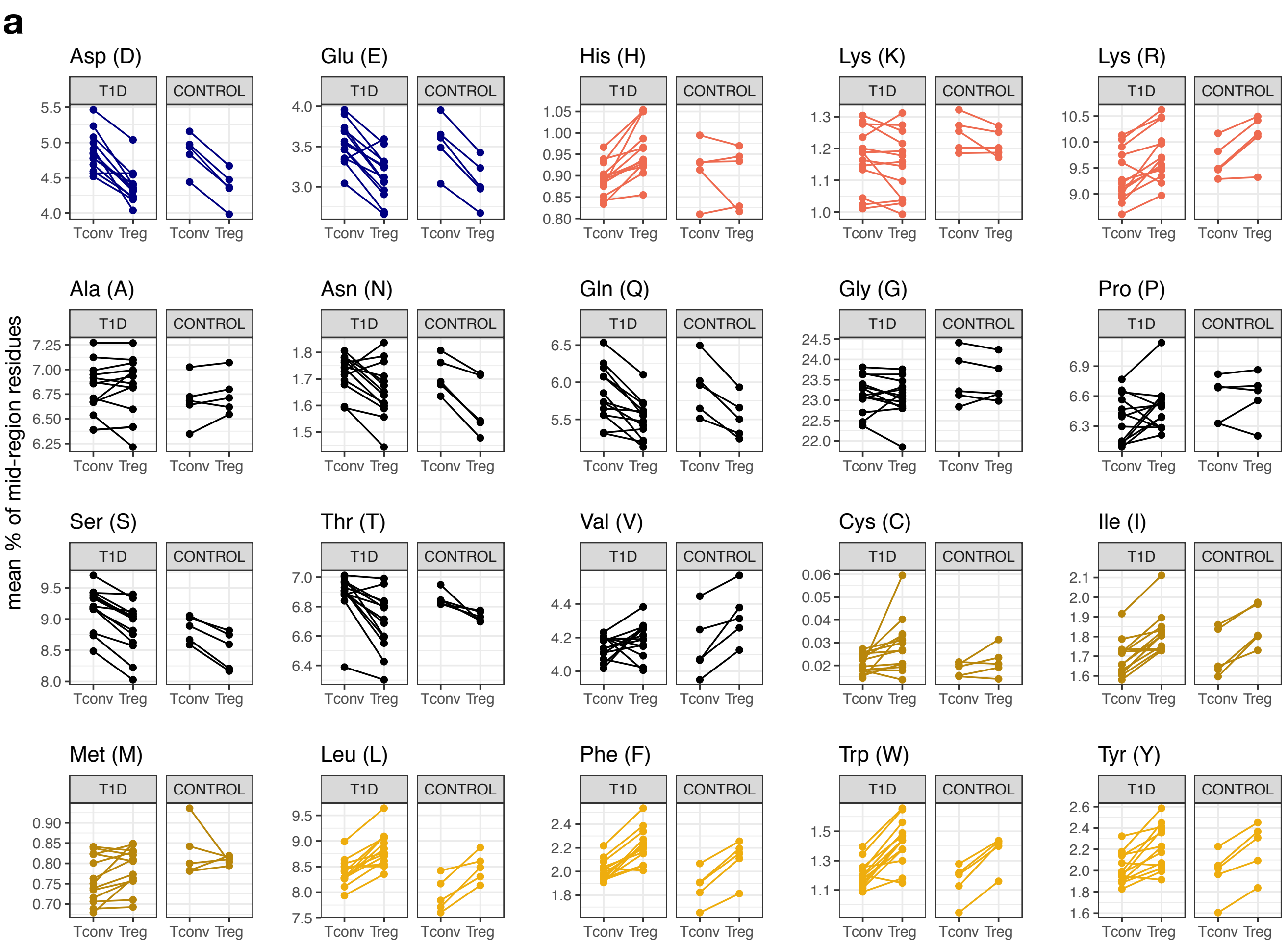


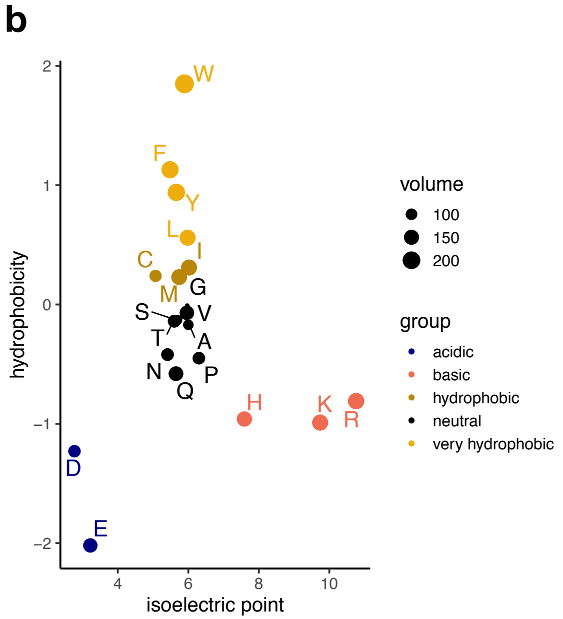


**Extended Data Figure 2. (a)** Mean percentage of each amino acid in each donor sample, separated by cell type and clinical phenotype and colored by amino acid groups as delineated in (b). T1D = "type 1 diabetes." **(b)** Categorization of amino acids by isoelectric point and interfacial hydrophobicity.

**Extended Data Figure 3.**

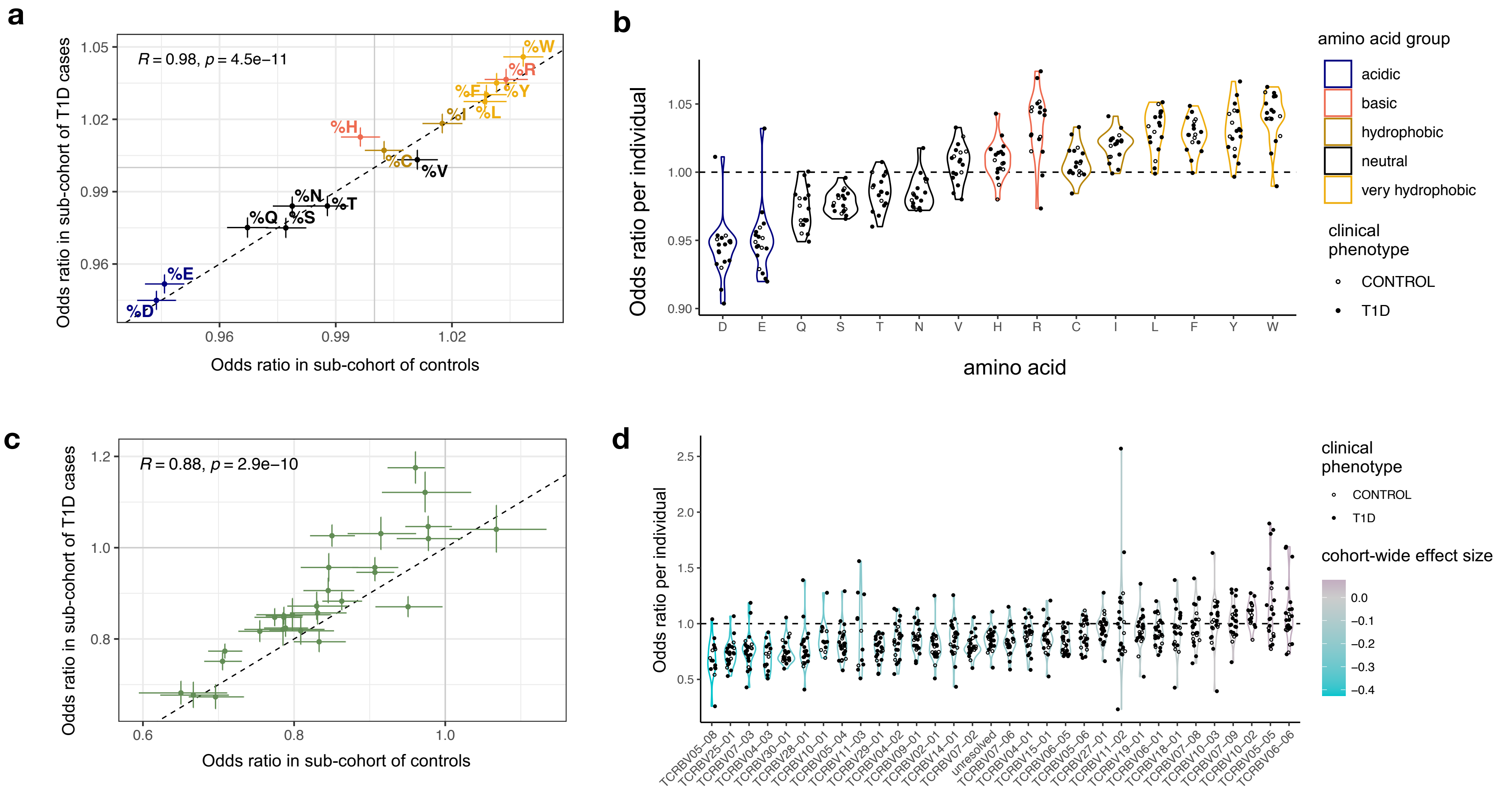

**Extended Data Figure 3. Consistency in TCR feature effect sizes across donors and clinical phenotypes.** **(a)** Treg odds ratio per standard deviation increase in CDR3βmr occupancy by each of the 14 relevant amino acids, estimated separately for the T1D cases in the discovery cohort (y axis) and the controls (x axis) **(b)** Treg odds ratio per standard deviation increase in CDR3βmr occupancy by each of the 15 relevant amino acids, estimated separately in each donor. **(c)** Treg odds ratio for the usage of each *TRBV* gene relative to the reference gene *TRBV05-01*, estimated separately for the T1D cases in the discovery cohort (y axis) and the controls (x axis) **(d)** Treg odds ratio for the usage of each *TRBV* gene relative to the reference gene *TRBV05-01*, estimated separately in each donor. *P* values in (a) and (c) are calculated by the t-test with Fischer transformation on Pearson's *R*.

### Extended Data Figure 4. Individualized thymic selection rates for *TRBV* and *TRBJ* genes

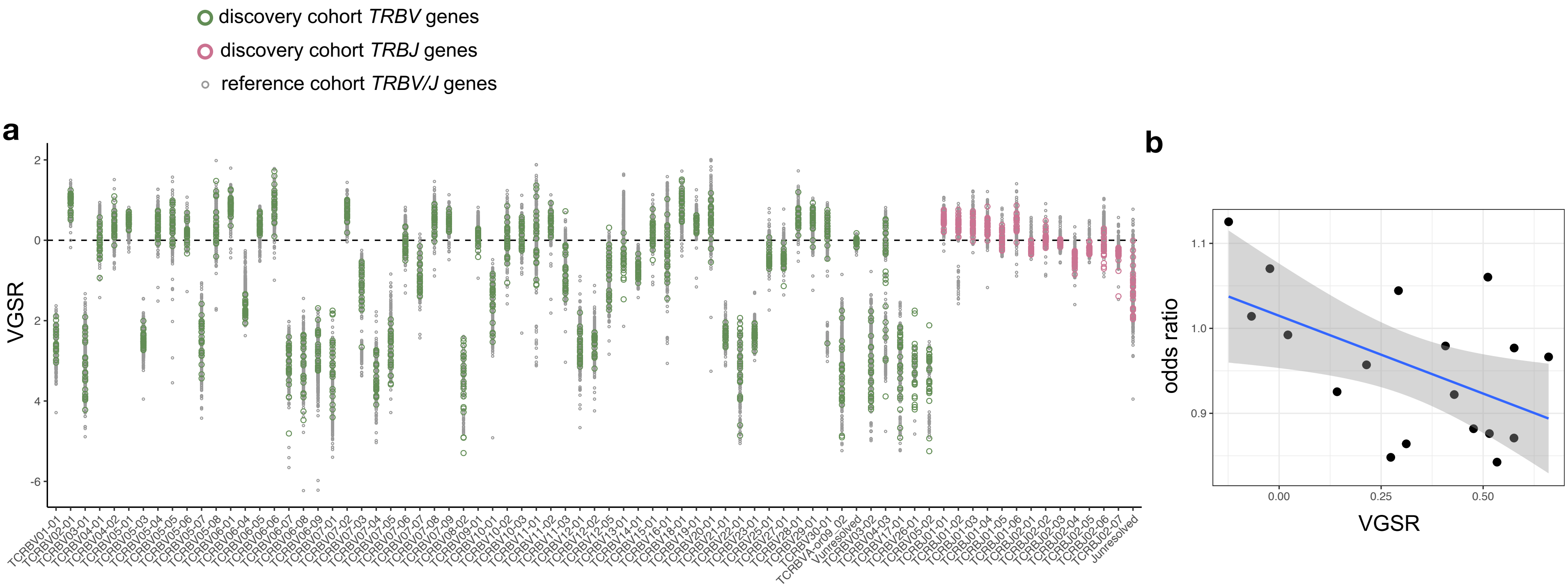


**Extended Data Figure 4. Individualized thymic selection rates for *TRBV* and *TRBJ* genes. (a)** V gene selection rates (VGSR) for each *TRBV* and *TRBJ* gene in each donor in the discovery cohort and in a reference cohort of 666 healthy donors, inferred by relative gene usage in productive reads versus nonproductive reads (**Supplementary Note**). **(b)** Individualized estimates for the effect of *TRBV19-01* (the most frequently used non-reference *TRBV* gene) on Treg fate exemplifies how the *TRBV* gene selection rate (VGSR) explains inter-individual variability in *TRBV* gene effects.

### Extended Data Figure 5.

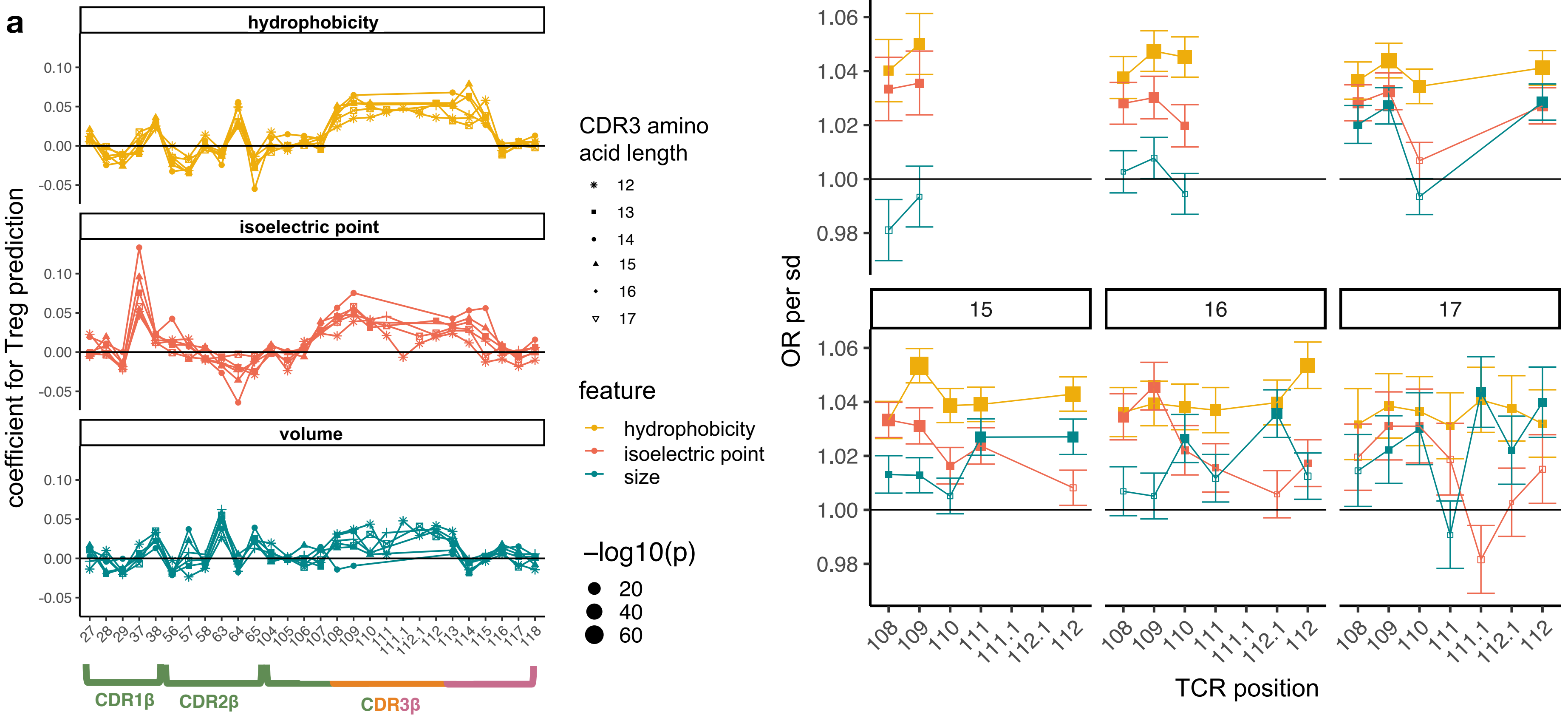


**Extended Data Figure 5. (a)** Estimated log odds ratio (per standard deviation) for each physicochemical feature at each CDRβ(1-3) loop position in each CDR3β length; features with an estimate > 0 are positively associated with Treg fate while features with an estimate < 0 are negatively associated. For each CDR3β length, all effects were estimated jointly in an L2-regularized logistic regression with 10-fold cross-validation (Methods). **(b)** Treg odds ratio per standard deviation increase in each physicochemical feature at each CDR3βmr position for each CDR3 length (Methods). Squares are sized inversely with the Wald test *P* values of the effect size they represent, filled in if this *P* value passes the Bonferroni significance threshold ($P < 0.05/81$ tests).

**Extended Data Figure 6. Cell type identification in single-cell data**

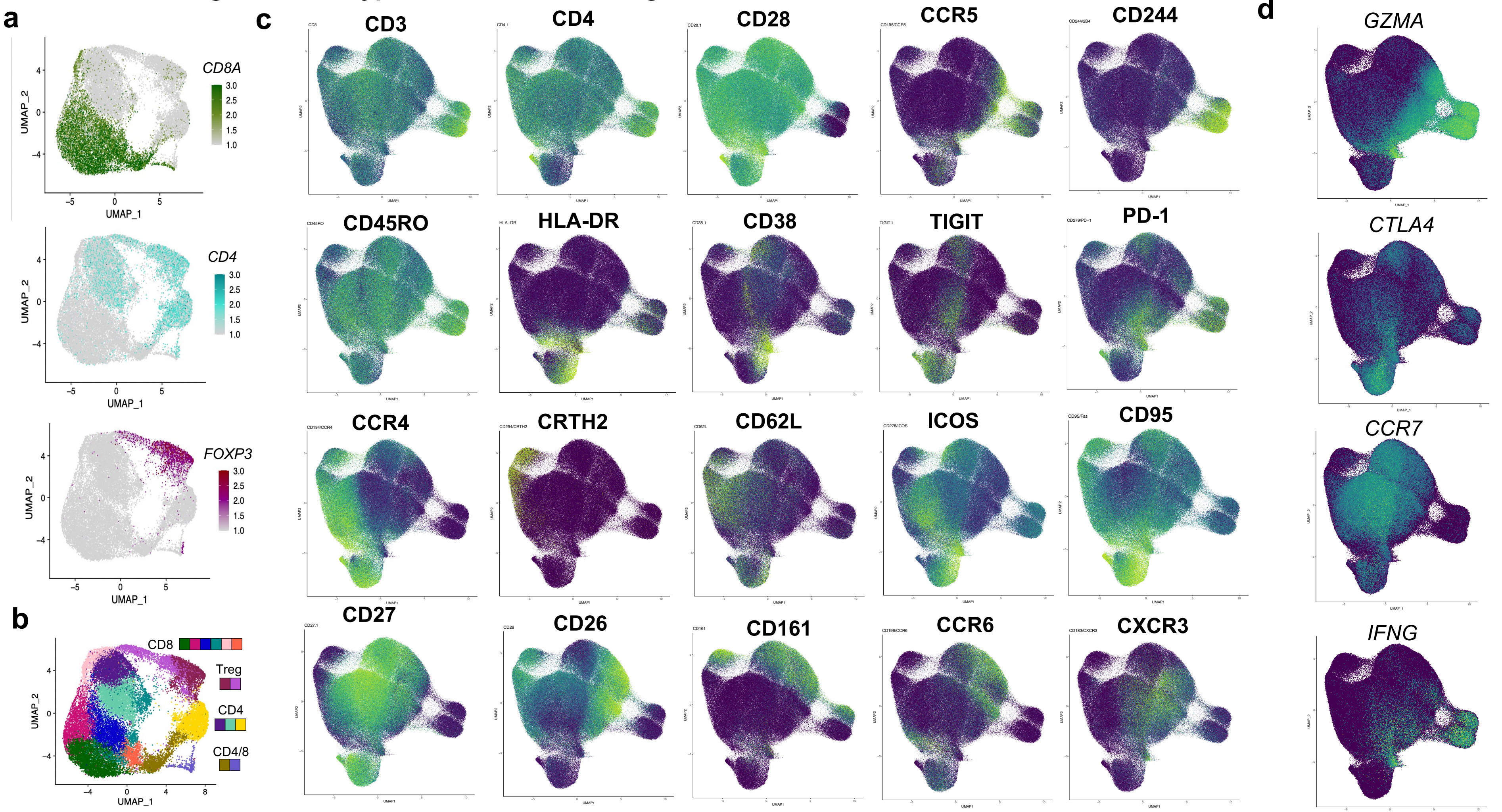

**Extended Data Figure 6. (a)** Log-normalized and scaled *CD8A, CD4* and *FOXP3* mRNA expression in T cells from breast tumor biopsies in Azizi et al. 2018, organized into a 2-dimensional embedding by Uniform Maniform Approximation and Projection (UMAP). **(b)** Louvain clustering of breast tumor microenvironment T cells. Broad cell type labels are indicated for each cluster in the surrounding legend. **(c)** Expression levels of key surface proteins measured by CITE-seq in the CD4+ reference single cell dataset[25] (low = purple, high = light green). Protein levels are normalized by the centered log-ratio (CLR) transformation (Methods). **(d)** Expression levels of key mRNA transcripts in the CD4+ reference single cell dataset[25] (low = purple, high = light green).

### Extended Data Figure 7. Single cell TiRP analysis details

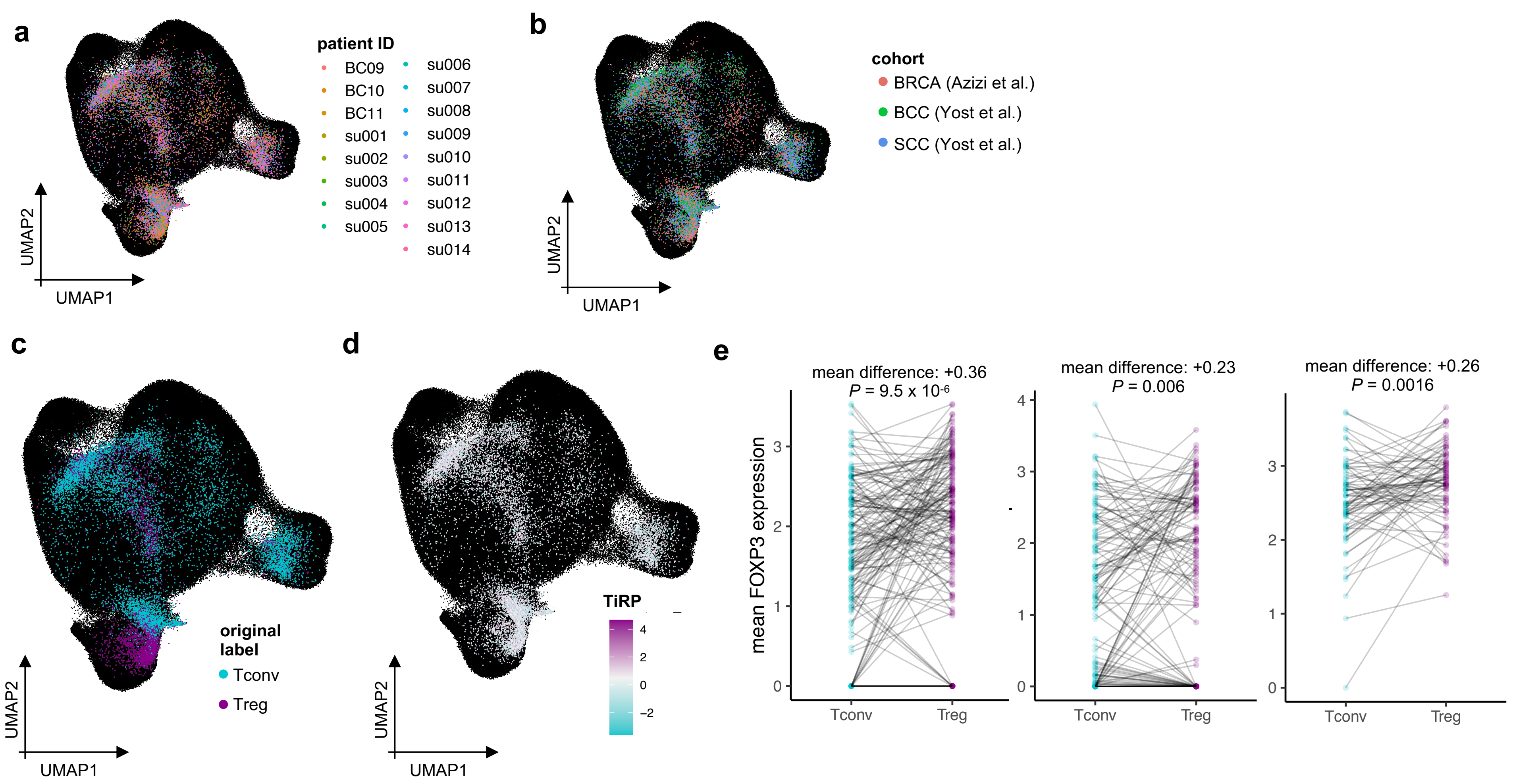

**Extended Data Figure 7. (a)** Tumor microenvironment T cells mapped into the reference embedding by Symphony, colored by donor to reveal successful integration of donors. **(b)** same as (a), colored by cancer type to reveal successful integration of cohorts. **(c)** Tumor microenvironment T cells mapped into the reference embedding by Symphony, colored by cell types derived from internal clustering (by Yost et al. for the SCC and BCC samples, and as depicted in Extended Data Figure 6 for the BRCA samples) to show the extent of concordance with Symphony's cell type solutions. **(d)** same as (a), colored by the TiRP score of their TCR. TiRP is scaled such that 0 corresponds to the mean score and one unit corresponds to one standard deviation of held-out bulk sequencing TCRs (Figure 5c). **(f)** *FOXP3* expression differences between Tregs and Tconvs within mixed clones of three representative donor samples. Each mixed clone is represented by a line connecting the average *FOXP3* expression of Tregs within the clone to the average *FOXP3* expression of Tconvs within the clone. Each *P* value is computed by a paired t-test comparing the mean *FOXP3* expression in Tregs to that in Tconvs within each mixed clone.

### Extended Data Figure 8.

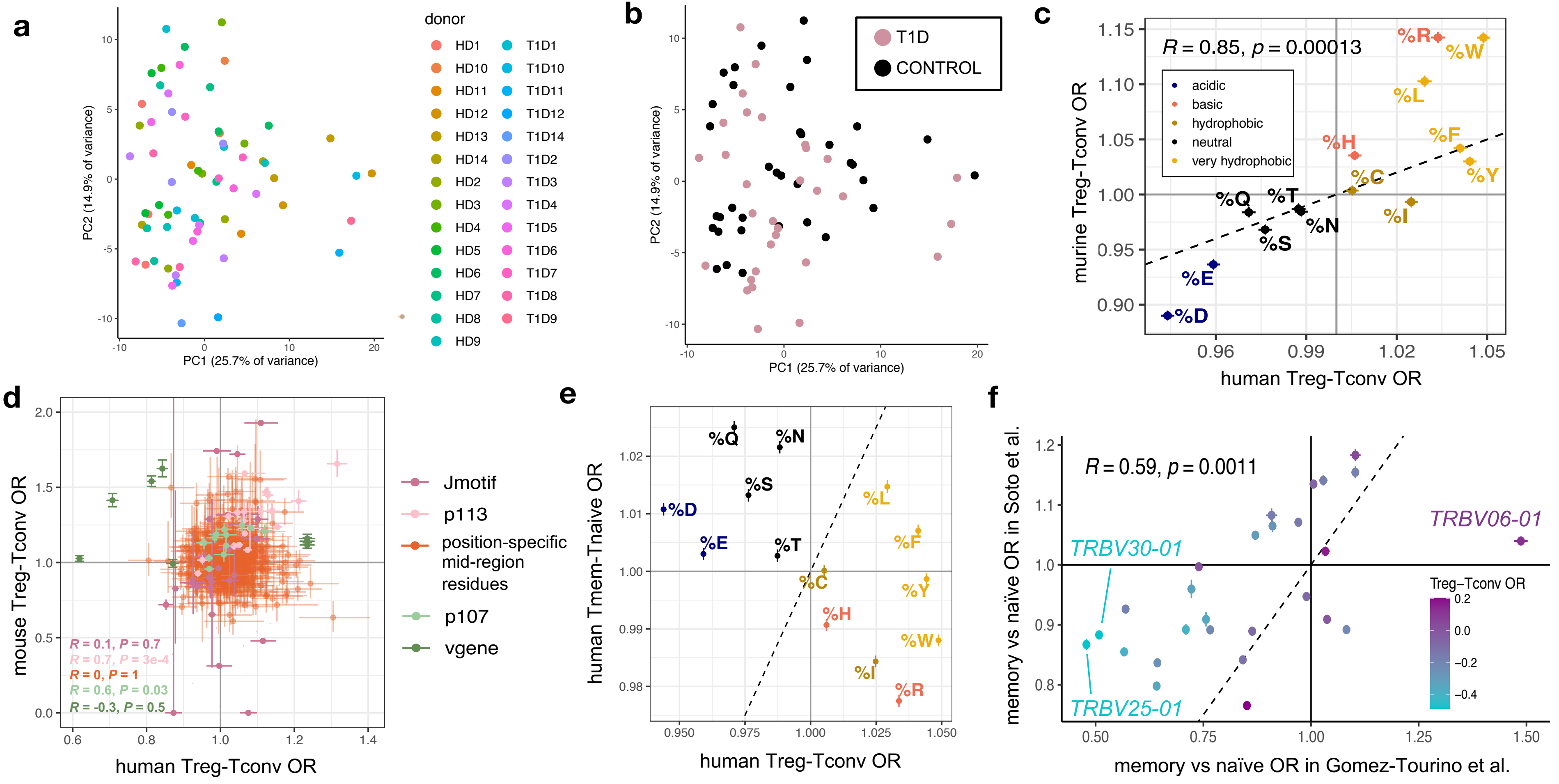

**Extended Data Figure 8. (a)** 67 samples from the replication cohort colored by donor ID and arranged by principal component space according to variation in TCR sequence feature frequencies. **(b)** Same as (a), colored by donor clinical phenotype. **(c)** Replication of CDR3βmr percent composition of amino acid effects in mice. Error bars correspond to 95% confidence intervals for ORs. **(d)** Lack of mouse-human correspondence for position-specific TCR feature effects. TCR features are colored by type; error bars denote OR 95% confidence intervals. Murine *TRBV* genes were mapped to their human homologs for comparison, only those with a human homolog are shown (Methods). **(e)** Overall lack of correspondence between Treg-Tconv OR and memory-naïve OR for CDR3βmr percent composition of amino acids. Error bars correspond to 95% confidence intervals, and amino acids are colored by the scheme in (c). **(f)** Replication of Tmemory-Tnaive *TRBV* gene odds ratios in an independent dataset of sorted memory and naïve T cells from 4 healthy donors[29]. *TRBV* genes are colored by their Treg-Tconv odds ratios. For (c), (d), and (f), *R* and *P* values are computed by a Student's t-test of Pearson's product-moment correlation. For (c)-(f), human Treg-Tconv OR result from meta-analysis across the discovery and replication cohorts.
