## Supplementary Note for "Repertoire analyses reveal TCR sequence features that influence T cell fate"

Multiple sources of inter-individual variation plausibly contribute to the thymic selection of *TRBV* genes. First, many of the 58 *TRBV* genes come in multiple allelic forms[1], and so the same *TRBV* gene in different individuals may have different functionality due to *TRBV* allelic variation. Second, because the *TRBV*-encoded region of the TCR is known to contact HLA molecules in the pMHC-TCR complex[2], genetic polymorphisms in the MHC locus likely influence the functionality and thymic selection of *TRBV* genes[3].

To examine inter-individual variation in the selection of *TRBV* genes, we analyzed nonproductive reads from each donor. Nonproductive reads are CDR3 sequences that are out of frame or contain a stop codon, which therefore cannot be expressed on the surface of the T cell. The DNA of the nonproductive TCR persists in the nucleus of the T cell after thymic selection, while the TCR from the homologous, successfully recombined allele is expressed on the cell surface. Because nonproductive TCRs never reach the cell surface, these data give a glimpse of *TRBV* and *TRBJ* usage prior to thymic selection. Several others[4–6] have successfully used nonproductive TCR reads to derive various individualized metrics of thymic selection. By focusing on the frequency of *TRBV* gene usage pre- and post- selection, our metric, “V gene selection rate” (VGSR), estimates the logarithm of the probability ratio *Q* in Elhanati et al.[5] with respect to individual *TRBV* genes.

For *TRBV* gene *i* in individual *j*,

$$\mathrm{VGS} = \ln\frac{(p_{ij}+1)/(p_j+1)}{(n_{ij}+1)/(n_j+1)}$$

where $p_{ij}$ is the number of productive reads with *TRBV* gene *i* in individual *j*, $p_j$ is the number of productive reads in individual *j*, $n_{ij}$ is the number of nonproductive reads with *TRBV* gene *i* in individual j, and $n_j$ is the number of nonproductive reads in individual *j*.

Thus, VGSR is the natural logarithm of the ratio of a *TRBV* gene’s frequency in the post-selection repertoire relative to the pre-selection repertoire, such that it is greater than 0 if and only if thymic selection acts to increase the repertoire frequency of the *TRBV* gene.

We computed VGSR for each *TRBV* gene in each donor of the discovery cohort as well as a reference cohort of 666 individuals, among the largest to date with deep TCR sequencing[7] (**Table 1**). Interestingly, donor-individualized VGSR estimates in the discovery cohort appeared to cluster around markedly different centroids for different *TRBV* genes. Though the reference cohort indeed revealed greater inter-individual variation, this still did not obscure the general differences between *TRBV* genes observed (**Extended Data Figure 4a**). In other words, inter-gene variance was greater than inter-individual variance, consistent with our hypothesis that *TRBV* genes' differential affinities to conserved sites of MHC may influence T cell fate in a consistent manner across individuals.

We then tested the effect of VGSR on the odds of Tregs fate using a mixed effects logistic regression model as described in the main text (**Methods**). When assessed as the only fixed covariate, VGSR was positively associated with Treg fate (OR = 1.01, 95% CI = 1.007 – 1.012, $P = 5.0 \times 10^{-15}$, LRT), indicating that Treg-associated *TRBV* alleles are overall rewarded by thymic selection. Considered in light of affinity-based Treg fate acquisition, this result is consistent with previous work showing that positive selection induces a greater change in the developing repertoire than negative selection, such that up to 95% of DP thymocytes die by neglect[8,9].

We next asked whether *TRBV* usage explained variance in Treg likelihood independent of thymic selection rate. We added VGSR as a covariate to our V-region mixed effects model, and tested whether *TRBV* gene identity would continue to explain a significant amount of variance in T cell fate. In this joint model, *TRBV* gene identity explained essentially the same amount of variance ($P = 7.9 \times 10^{-866}$, LRT, 30.9% of total variance explained by the TCR compared to 30.8%). These results indicate that *TRBV* usage does exert an influence on Treg fate that is independent of all inter-individual variance captured by VGSR, including *TRBV* allelic variation and MHC polymorphisms. In this joint model, VGSR was negatively associated with Treg fate (OR = 0.965, 95% CI = 0.956 – 0.975, $P = 8.6 \times 10^{-13}$), suggesting that allelic variation influences negative selection, but that this effect is minor compared to overall differences in the positive selection of *TRBV* genes.

To further confirm that the cohort-wide *TRBV* gene effect estimates were robust to inter-individual variation, we estimated *TRBV* gene effects in each donor separately. The ordering of *TRBV* genes in each donor generally followed cohort-wide estimates, and VGSR indeed helped to account for inter-individual differences (**Extended Data Figure 4b**). Thus, we proceeded with the cohort-wide *TRBV* gene effect sizes adjusted for inter-individual variation in *TRBV* thymic selection by the VGSR covariate. These adjusted effect sizes are used in the computation of Treg-intrinsic regulatory potential (TiRP, **Supplementary Table 7**), such that TiRP can be applied to new TCR data without any information regarding nonproductive reads, HLA haplotypes, or *TRBV* alleles. Without this information, the resultant score conveys the TCR-intrinsic regulatory potential of the given TCR with a presumed average VGSR. Given the limited amount of variance explained by VGSR (<1% of the total variance explained by the TCR, **Figure 3c**), TiRP calculation without these data adequately approximates the regulatory potential of the TCR.

As expected from the absence of J-region-MHC structural contacts, the analogous calculation for *TRBJ* gene selection rates revealed minimal inter-individual variance (**Extended Data Figure 4a**) and did not have a significant association with Treg fate.
